# Molecular insights into the Darwin paradox of coral reefs from the sea anemone Aiptasia

**DOI:** 10.1101/2022.10.02.510507

**Authors:** Guoxin Cui, Migle K. Konciute, Lorraine Ling, Luke Esau, Jean-Baptiste Raina, Baoda Han, Octavio R. Salazar, Jason S. Presnell, Nils Rädecker, Huawen Zhong, Jessica Menzies, Phillip A. Cleves, Yi Jin Liew, Cory J. Krediet, Val Sawiccy, Maha J. Cziesielski, Paul Guagliardo, Jeremy Bougoure, Mathieu Pernice, Heribert Hirt, Christian R. Voolstra, Virginia M. Weis, John R. Pringle, Manuel Aranda

## Abstract

Symbiotic cnidarians such as corals and anemones form highly productive and biodiverse coral-reef ecosystems in nutrient-poor ocean environments, a phenomenon known as Darwin’s Paradox. Resolving this paradox requires elucidating the molecular bases of efficient nutrient distribution and recycling in the cnidarian-dinoflagellate symbiosis. Using the sea anemone Aiptasia, we show that during symbiosis, the increased availability of glucose and the presence of the algae jointly induce the coordinated upregulation and re-localization of glucose and ammonium transporters. These molecular responses are critical to support symbiont functioning and organism-wide nitrogen assimilation through GS/GOGAT-mediated amino-acid biosynthesis. Our results reveal crucial aspects of the molecular mechanisms underlying nitrogen conservation and recycling in these organisms that allow them to thrive in the nitrogen-poor ocean environments.

**One-sentence summary:** Whole-organism nitrogen assimilation fueled by glucose from symbiotic algae enables corals to flourish in oligotrophic waters.

---

The ability of corals to build one of the planet’s most biodiverse and productive ecosystems in the nutrient-poor seawater of the subtropics and tropics, often referred to as “ocean deserts”, has both fascinated and puzzled scientists since it was first noted by Darwin (*1*). The foundation of these ecosystems is the symbiotic relationship between cnidarian host animals and photosynthetic dinoflagellate algae (*2, 3*) of the family Symbiodiniaceae (*4*), which live in specialized vacuoles (known as “symbiosomes”) inside the gastrodermal cells that line the gastric cavity of the host. The hosts and algae, together with a diverse assemblage of microorganisms, form metaorganisms known as holobionts (*5*). Algal photosynthesis provides fixed organic carbon for energy and biosynthesis and covers the majority of the hosts’ energy demands (*2, 3*). However, the provision of organic carbon is not the only important function attributed to the algal endosymbionts. Nitrogen is one of the primary growth-limiting nutrients in coral-reef ecosystems (*2, 6*), and the algae have been thought to be the main contributors to nitrogen acquisition and recycling (*7–11*) due to their high capacity for ammonium assimilation (*12*). However, some evidence has also suggested an active role for the host in nitrogen assimilation (*13*), a view supported by the recent realization that the host also possesses the enzymatic machinery to recycle ammonium via the glutamine synthetase / glutamate synthase (GS/GOGAT) system (*14–16*).

The importance of photosynthetically fixed carbon and of nitrogen assimilation and conservation for the ecological success and productivity of these metaorganisms is well established. However, we still do not know how fixed carbon is moved from the algae to the various host cells as well as the respective contribution of the host and algae to nitrogen assimilation and conservation. Unraveling these mechanisms is critical for our understanding of holobiont functioning and ecological productivity. In this study, we used the sea anemone Aiptasia, which, like corals, harbors symbiotic dinoflagellates in the family Symbiodiniaceae (*17–19*), to investigate these matters in detail in an experimentally tractable model system that has the advantage of allowing comparisons between symbiotic and non-symbiotic individuals.

## Results

### Modulation of gene expression by tissue type and symbiotic state

To investigate nutrient fluxes within the cnidarian-algal symbiosis in the context of the spatial organization of the holobiont, we first isolated gastrodermal and epidermal tissues from both symbiotic and aposymbiotic anemones (thus, four tissue types in total) using laser microdissection (LMD; Fig. S1) and analyzed their transcriptomic profiles via RNA-Seq (Data S1). A principle-component analysis (PCA) showed that samples clustered by both tissue layer (PC1: ~24% of the variance) and symbiotic state (PC2: ~18% of the variance) (Fig. 1A). Symbiosis induced extensive changes in gastrodermal gene expression as well as significant (although more limited) changes in epidermal gene expression (Fig. 1B). Using a multi-factorial differential-expression analysis including both tissue identity and symbiotic state, we identified 5,414 gene-expression signatures linked to one or both of these factors. Hierarchical clustering of these genes identified five modules showing distinct expression patterns (Fig. 1C; Data S2): genes in modules 1 and 2 were symbiosis-induced and -repressed, respectively; genes in module 3 were symbiosis-induced and gastrodermis-specific; and genes in modules 4 and 5 were gastrodermis- and epidermis-specific, respectively.

**Fig. 1.**
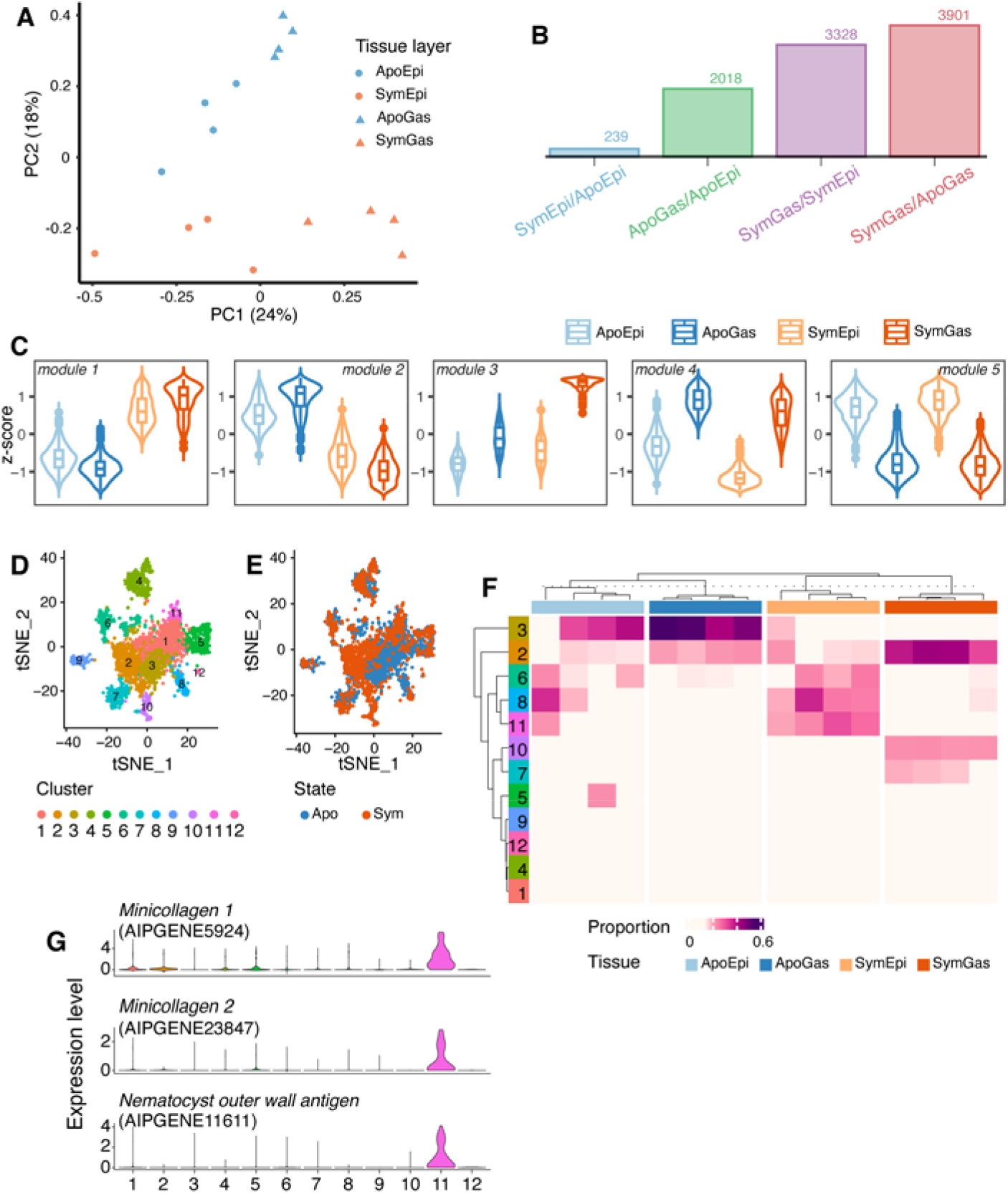
Transcriptomic profiles of tissues and cells isolated from symbiotic and aposymbiotic Aiptasia. (A) Principal-component analysis of laser-microdissection RNA-Seq data generated from four biological replicates of each of four tissue types. Apo, aposymbiotic; Sym, symbiotic; Gas, gastrodermis; Epi, epidermis. (B) The numbers of differentially expressed genes (DEGs) identified in the pairwise comparisons (*q* value < 0.05). (C) The five modules determined by hierarchical-cluster analysis on the expression levels (scaled z-scores) of the DEGs. (D) Clustering of 2,698 Aiptasia cells into 12 clusters (each a different color) in *t*-SNE space based on their transcription profiles. Each dot represents one cell. (E) The same 12 clusters with the symbiotic state of the source animal indicated for each cell. (F) The proportions of the 12 cell clusters in the bulk-sequenced tissue-specific samples (four samples per tissue type). The cellular complexities of the tissue samples were characterized by MuSiC. (G) The expression patterns of three Cluster 11 marker genes across all cells.

We next performed functional-enrichment analyses to identify the predominant functional categories of the genes within the modules (*p* < 0.05; Fig. S2). Module 1 was enriched for genes associated with both carbon (Table S1) and nitrogen (Table S2) cycling (including at least one predicted glucose transporter and one predicted ammonium transporter) as well as protein post-translational modifications (Table S3), presumably reflecting the metabolic changes induced by symbiosis in both tissue layers. Module 2 was enriched for genes involved in food digestion (Table S4), presumably reflecting the dependence of aposymbiotic animals on heterotrophic food sources. Not surprisingly, Module 3 (symbiosis- and gastrodermis-specific) was enriched for functions related to organization of the symbiosome (Table S5) and the transport of various key metabolites, such as glucose (Table S6) and cholesterol (Table S7), whereas the gastrodermis-specific Module 4 included many digestion-related genes (Table S8), and the epidermis-specific Module 5 was enriched for genes involved in cnidocyte function (Table S9) and responses to mechanical stimuli (Table S10).

Given the central importance of the GS/GOGAT cycle in ammonium assimilation and the strong symbiosis-induced upregulation of the Aiptasia genes for both enzymes at the whole-organism level (Fig. S3A) (*8, 14–16*), we were surprised that these genes did not appear in either Module 1 or Module 3. However, as we also did not find any appreciable difference between gastrodermis and epidermis in the expression levels of either GS or GOGAT (Fig. S3B), the most parsimonious interpretation is that the tissue-specific-expression data are somehow misleading in this case (see Discussion) and that GS and GOGAT in fact participate in enhanced ammonium assimilation by both major tissues of symbiotic anemones, consistent with the transporter-localization and NanoSIMS data presented below.

### Cell-type-specific responses to symbiosis

Although many of the DEGs identified in the analysis of tissue-specific expression had been reported previously to be symbiosis-regulated (*14, 15, 20*), our analysis began to provide the spatial resolution needed to investigate their functions further. To gain higher-resolution spatial information, we next performed single-cell RNA-Seq on isolated cells using the 10x Genomics platform. We retrieved gene-expression information from 2,698 cells, of which 1,453 originated from aposymbiotic and 1,245 from symbiotic anemones. Following *t*-distributed stochastic-neighbor-embedding (*t*-SNE) analysis, we grouped the cells into 12 clusters with distinct gene-expression profiles (Fig. 1D) and identified potential cell-type-marker genes (adjusted *p* < 0.01 and average fold-change > 2; Data S3). Most of these clusters were shared between symbiotic and aposymbiotic animals, but some clusters were largely specific to one symbiotic state (Fig. 1E, Fig. S4).

To investigate the tissue origins of the cells in the 12 clusters, we integrated our single-cell and tissue-specific data by performing a deconvolution analysis using MuSiC (*21*), which indicated that five of the cell clusters (1, 4, 5, 9, 12) were present at low abundance (< 10% of the total cells) in all of the tissue samples (Fig. 1F; see Discussion). The other seven clusters exhibited tissue- and/or symbiotic-state-specific associations (Fig. 1F). Cluster 11 was present exclusively in the epidermis, and its highly expressed marker genes are cnidocyte-specific, as identified in other cnidarian species (Fig. 1G, Table S11) (*22–24*), so that this cluster appears to represent cnidocytes. In contrast, Clusters 2, 7, and 10 were highly represented in the gastrodermal samples, with the latter two groups being specific to symbiotic animals (Fig. 1F). Many of the highly expressed marker genes for these clusters (Table S12) had been identified previously as displaying gastrodermis-specific expression in the sea anemone Nematostella (*25*), the stony coral Stylophora (*26*), and/or the soft coral Xenia (*23*), consistent with a gastrodermal origin for these cells.

We then further examined the functions of the putative symbiotic gastrodermal cells through a gene-set-enrichment analysis of their marker genes (Data S4 and S5). Cluster 10 marker genes were enriched for ones associated with lysosomal organization and function (Table S13), glucose transport and its positive regulation (Table S14), and cholesterol homeostasis (Table S15), whereas Cluster 7 marker genes were enriched for functions associated with the extracellular matrix, cell adhesion, and cell-cell signaling (Table S16). Thus, cluster 10 appears to be symbiotic cells, whereas cluster 7 may contain gastrodermal cells that are free of algal cells.

### Symbiosis induces elevated expression and relocalization of glucose and ammonium transporters

To explore further the tissue- and cell-specific incorporation of carbon and nitrogen, we focused on the major glucose and ammonium transporters. At least six putative glucose transporters and two putative ammonium transporters showed significant changes in tissue- and/or cell-specific expression in response to symbiosis (Fig. 2, A and B; Tables S1, S2, S6, and S17), so we examined the functions and localizations of five of these transporters in more detail.

**Fig. 2.**
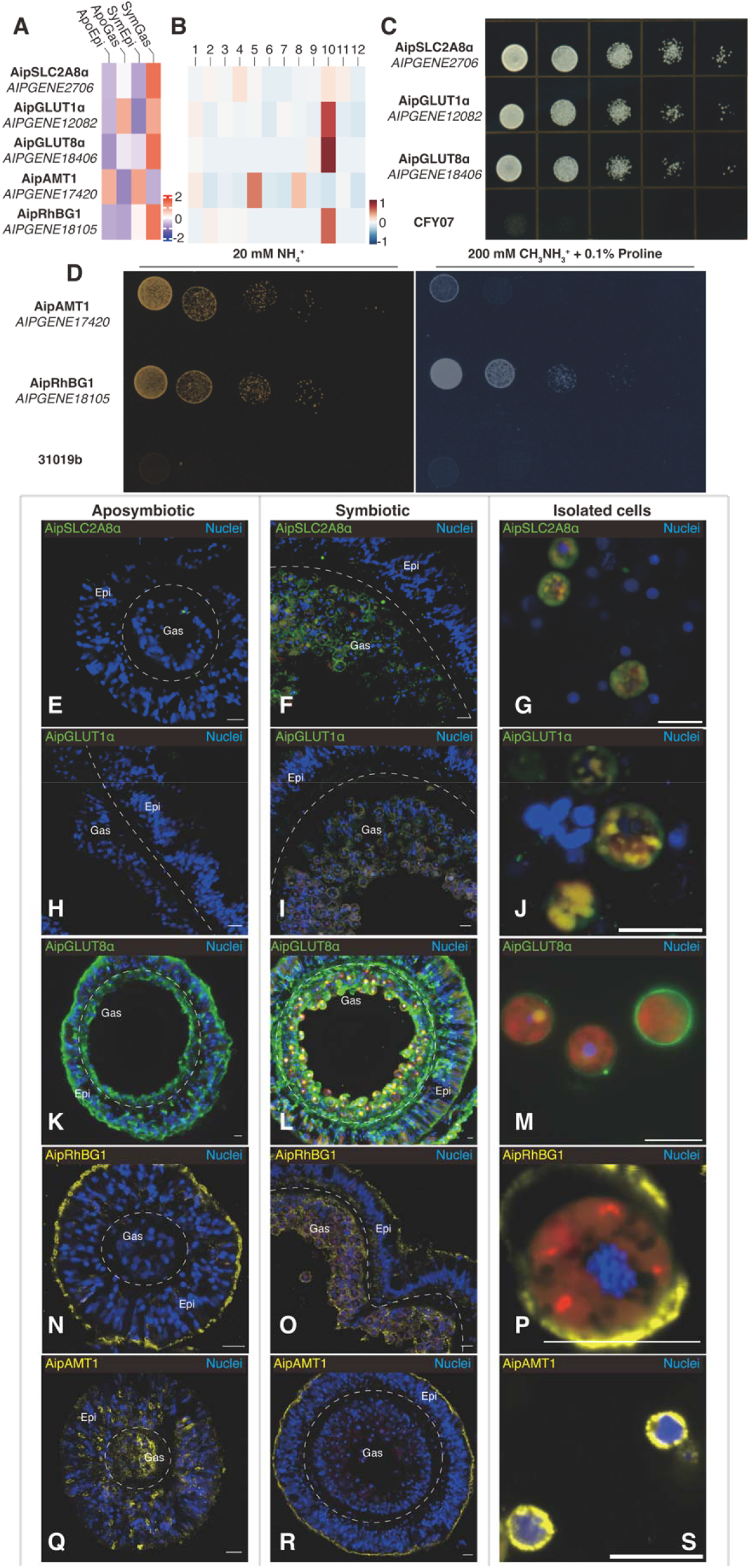
Altered expression and relocalization of glucose and ammonium transporters during symbiosis. (A, B) Expression patterns of mRNAs for glucose and ammonium transporters at the tissue (A) and cell (B) levels. (C) Rescue experiments on yeast mutant CFY07 (lacking all sugar transporters; see Materials and Methods) expressing one of the putative Aiptasia glucose transporters and spotted in a dilution series on medium containing glucose as the sole carbon source. (D) Rescue experiments on yeast mutant strain 31019b (lacking all ammonium transporters; see Materials and Methods) expressing Aiptasia AipAMT1 or AipRhGB1. The plate on the left contained 20 mM ammonium as the sole nitrogen source. The plate on the right contained 0.1% proline as a nitrogen source plus 200 mM of the toxic ammonium analogue methylammonium. (E – S) Immunofluorescence staining of glucose (E – M) and ammonium (N – S) transporters in tissue sections of aposymbiotic (E, H, K, N, Q) or symbiotic (F, I, L, O, R) anemones, as well as in cells isolated from symbiotic animals (G, J, M, P, S). Scale bars (all panels), 10 μm.

First, we conducted rescue experiments in yeast mutants to test the putative transporter activities of the gene products. Indeed, each one rescued the appropriate yeast mutant (*27, 28*), allowing its growth on the relevant selective medium (Fig. 2C; Fig. 2D, left), verifying that each indeed had the predicted glucose- or ammonium-transporter activity.

We next generated antibodies specific for each of these five transporters (see Materials and Methods; Fig. S5; Table S18) and used these antibodies to examine their localizations by immunofluorescence staining. AipSLC2A8α (Fig. 2, E – G; Fig. S6, A – C) and AipGLUT1α (Fig. 2, H – J; Fig. S6, D – F) were detected primarily or exclusively in symbiotic gastrodermis as well as in isolated symbiotic cells. For AipGLUT1α (*AIPGENE12082*), the apparent discrepancy between these results and those on tissue-specific transcript levels (Fig. 2A; Table S17) may reflect a misleading feature of the latter, given that *AIPGENE12082* transcript levels were substantially upregulated in the putatively gastrodermal Cluster 10 cells in the single-cell analysis (Fig. 2B; Table S14; see Discussion). Their localization patterns suggest that these transporters might serve mainly to move photosynthetically produced glucose (*29*) across the symbiosome membrane into the host-cell cytoplasm and/or from the symbiotic cells to the rest of the organism. In contrast, AipGLUT8α, despite its high apparent tissue (Fig. 2A; Tables S6 and S17) and cell (Fig. 2B) specificity as seen by RNA-Seq, was detected both in the outer (seawater-facing) surface of the epidermis and the inner (body-cavity-facing) surface of the gastrodermis in aposymbiotic anemones (Fig. 2K; Fig. S6G), suggesting that in such animals it might have a role in the scavenging of environmental glucose. In symbiotic anemones, however, this localization was largely replaced by one in which the peripheries of the gastrodermal cells, or perhaps their symbiosomes (the images do not have sufficient resolution to tell), were heavily stained, along with substantial staining of the gastrodermis-epidermis boundary (Fig. 2, L and M; Fig. S6, H and I), suggesting a role for AipGLUT8α also in the dissemination of photosynthetically produced glucose.

Localization of the two ammonium transporters also changed in response to symbiosis. AipRhBG1 was observed primarily at the outer surface of the epidermal cells in aposymbiotic animals (Fig. 2N; Fig. S6J), suggesting a role in the excretion of excess ammonium in the heterotrophic animals. In contrast, in symbiotic animals, although the protein was still observed in the outer layer of the epidermis, it was now most prominent around the gastrodermal cells and along the gastrodermis-epidermis boundary (Fig. 2, O and P; Fig. S6, K and L), suggesting that under these conditions, it functions in the uptake of ammonium for both animal and algal use. Consistent with the hypothesis that AipRhBG1 can transport ammonium both out of and into Aiptasia cells, yeast cells expressing AipRhBG1 as their sole ammonium transporter were relatively resistant to the toxic ammonium analog methylammonium (Fig. 2D, right panel), suggesting that this compound did not accumulate to high levels in the cells (*30*). In contrast, yeast cells expressing only AipAMT1 were much more sensitive to the drug (Fig. 2D, right panel), suggesting that this transporter functions only in ammonium uptake. Consistent with this hypothesis, immunofluorescence staining found AipAMT1 diffusely localized in aposymbiotic animals (Fig. 2Q; Fig. S6O) but concentrated in the outer layer of the epidermis in symbiotic animals (Fig. 2, R and S; Fig. S6, P and Q).

### Effects of metabolism and symbiosis on gene expression and protein localization

We next asked whether the symbiosis-specific changes in gene expression and transporter localization were triggered simply by the increased availability of glucose when algae are present or by some other aspect of algal presence. The data presented above suggested that both photosynthetically derived glucose and ammonium are distributed throughout the symbiotic animals, which might, in turn, induce a high expression of the central GS/GOGAT ammonium-assimilation machinery in both epidermal and gastrodermal cells. To test this hypothesis, we provided aposymbiotic anemones with supplemental glucose and analyzed gene-expression changes in comparison to aposymbiotic and symbiotic anemones without added glucose. We found that glucose treatment indeed significantly altered the whole-organism transcriptomic profiles (Fig. 3A). In particular, the expression levels of the AipAMT1 ammonium transporter and of several key nitrogen-metabolism enzymes, including GS and GOGAT, were essentially the same in symbiotic and glucose-treated aposymbiotic animals (Fig. 3B). Thus, some aspects of a shift toward organism-wide assimilation of ammonium in symbiotic anemones appear to be a direct response to the increased availability of glucose.

**Fig. 3.**
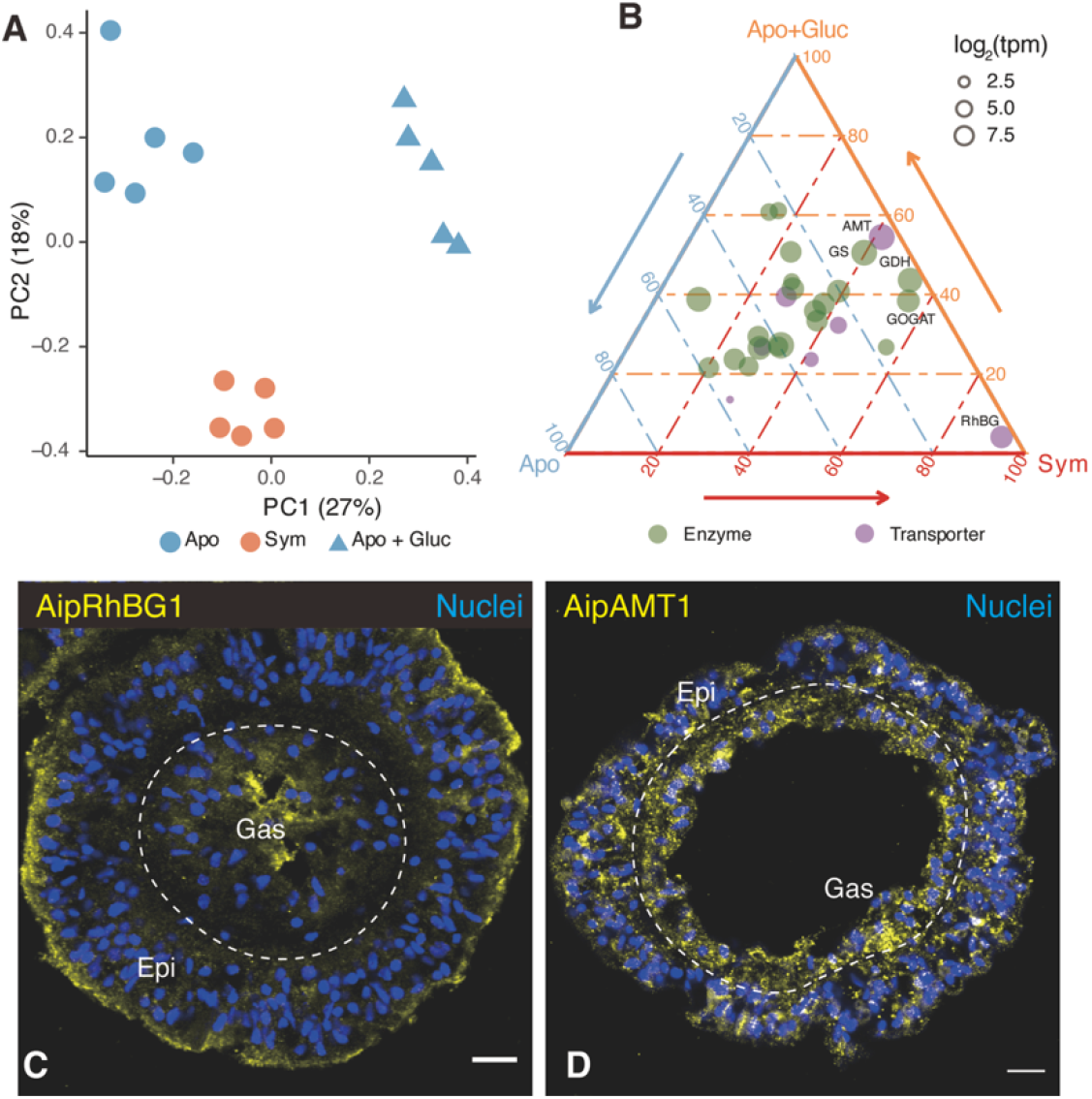
Expression changes of nitrogen-metabolism genes in response to glucose supplementation. (A) Principal-component analysis of whole-animal Aiptasia RNA-Seq data from an experiment comparing expression in aposymbiotic anemones with added glucose (Apo + Gluc) to that in aposymbiotic and symbiotic anemones without added glucose. (B) Ternary plot showing relative expression levels of genes associated with “nitrogen-metabolism pathway” (KEGG ko00910) in Aiptasia. Each dot represents a gene with coordinates representing the relative proportion of its expression level in each condition relative to its overall expression across the three conditions. (C, D) Immunofluorescence staining of AipRhGB1 (C) and AipAMT1 (D) in aposymbiotic Aiptasia supplied with glucose. Scale bars (all panels), 10 μm.

However, other observations suggest a more complicated picture. First, AipRhGB1 mRNA levels were not increased in glucose-treated aposymbiotic anemones (Fig. 3B). Moreover, immunofluorescence staining of AipRhGB1 and AipAMT1 in the glucose-treated aposymbiotic anemones showed no obvious change in the localization of either protein (Fig. 3, C and D; Fig. S7; cf. Fig. 2, N and Q). Thus, both the symbiosis-induced increase in AipRhBG1 expression and the changes in the localization of both ammonium transporters (see above) appear to depend on the actual presence of the algae. Thus, although the metabolic response to assimilate ammonium via the GS/GOGAT pathway appears to depend only on the increased availability of glucose, a separate, algae-dependent mechanism appears to exist for the regulation of nitrogen provision to the algae.

### Coordinated incorporation of carbon and nitrogen in both gastrodermis and epidermis of symbiotic animals

The data presented above suggest that symbiosis induces the organism-wide distribution of photosynthetically produced glucose and the assimilation of ammonium. To test this idea further, we incubated animals with ^13^C bicarbonate and ^15^N ammonium and quantified the assimilation of carbon and nitrogen in the gastrodermis and epidermis using nanoscale-secondary-ion mass spectrometry (NanoSIMS). The ^13^C-labeled glucose should provide both gastrodermal and epidermal cells with both the ATP and the carbon backbones required for the assimilation of ammonium into amino acids, nucleic-acid bases, and other compounds. Thus, we expected a strong spatial correlation of the ^13^C and ^15^N signals in both major host tissues in symbiotic animals. As expected, the assimilation of both ^13^C and ^15^N was low in both tissues of aposymbiotic animals (Fig. 4, A and B), and ^13^C and ^15^N assimilation were not well correlated (Fig. 4E). In contrast, symbiotic anemones incorporated significantly more of both isotopes in both major tissue layers (Fig. 4, C and D). Importantly, we also found a strong and statistically significant spatial correlation of the ^13^C and ^15^N signals in symbiotic anemones (at a spatial resolution of 88 nm: Fig. 4E, Fig. S8), suggesting that these elements might be co-incorporated into similar biosynthetic products in both major tissue layers.

**Fig. 4.**
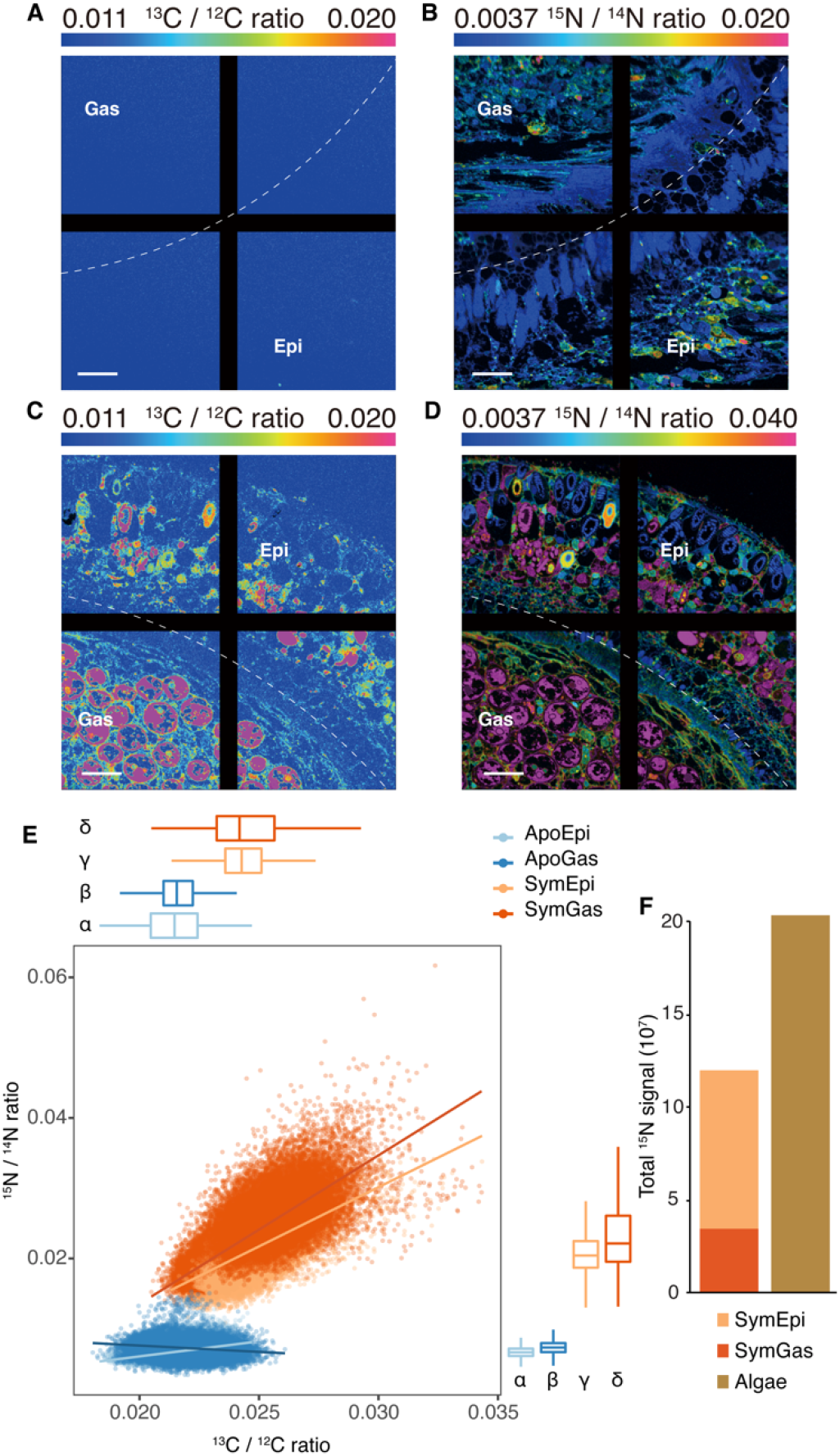
Carbon and nitrogen incorporation into gastrodermis and epidermis in aposymbiotic and symbiotic Aiptasia. (A - D) Representative images showing the distributions of ^13^C/^12^C (A, C) and ^15^N/^14^N (B, D) ratios in aposymbiotic (A, B) and symbiotic (C, D) Aiptasia. The ratios are displayed as Hue Saturation Intensities in which blue indicates the natural-abundance isotope ratio, with a shift toward magenta indicating an increasing ^13^C or ^15^N incorporation level. Scale bars, 10 μm. The black lines are a feature of the NanoSIMS display without biological significance. (E) The correlations between ^13^C/^12^C and ^15^N/^14^N ratios in epidermis and gastrodermis from aposymbiotic and symbiotic anemones. ^13^C/^12^C and ^15^N/^14^N ratios were quantified at the pixel level (spatial resolution, 88 nm) for regions of interest (see Fig. S8) across whole tentacle sections. Dots in the scatter plot represent the individual bins calculated, and the trendlines were estimated based on a generalized linear model. Box-and-whisker plots show the distributions of ^13^C/^12^C and ^15^N/^14^N ratios for each tissue layer; Greek letters indicate statistically significant (*p* < 0.001) differences between tissue layers as calculated using one-way ANOVA with Games-Howell post hoc tests. (F) Total absolute ^15^N signal in animal tissue layers and algal cells of symbiotic anemones. The marking method shown in Fig. S8 was used to separate animal tissue and algal cells for each section. The total ^15^N signal was then calculated by summing raw pixel values collected from the ^15^N channel as shown in Fig. S8B.

To further assess the contribution of ammonium assimilation by the host tissue layers to the overall nitrogen assimilation by the holobiont, we extracted absolute ^15^N signals from animal tissues and algal cells, respectively (marking method: Fig. S8B). The host tissues (excluding algae) contained 35 to 41% of the total ^15^N signal in symbiotic anemones (Fig. 4F), and no statistically significant difference for the absolute ^15^N signals was found between host and algae (paired *t*-test, *p* = 0.41). Remarkably, the epidermis actually incorporated significantly more ^15^N than the corresponding gastrodermis (paired *t*-test, *p* = 0.02). Thus, both gastrodermis and epidermis of the host contribute substantially – on a par with the algae – to overall holobiont nitrogen incorporation.

## Discussion

For a century and a half after it was first noted by Darwin (*1*), the apparent paradox of highly productive and species-rich coral reefs thriving in nutrient-poor ocean waters remained a mystery. The first major step in resolving this paradox was the discovery that most of the organic carbon required by the coral animals for energy and biosynthesis is provided through photosynthesis by their endosymbiotic dinoflagellate algae (*2, 3*). A second major step was the recognition that the problem of limiting nitrogen in marine environments (*2, 6*) was solved, in part, by nitrogen conservation and/or recycling within the coral holobiont (*8, 11*). However, these pioneering studies left many critical questions unanswered. In what form, and how, is the photosynthetically fixed carbon passed from the algae to the host gastrodermal cells and then distributed to the other host cells and tissues? How is excess nitrogen (when present) disposed of, and how is available nitrogen acquired from the environment when needed? What are the respective roles of the animal and its algal partner in the conservation and/or recycling of nitrogen within the holobiont? And what aspect(s) of algal presence trigger the adjustments in gene expression and metabolism that the host must make for an effective symbiosis? In this study, we addressed these questions through a combination of whole-animal, tissue-specific, and single-cell analyses of gene-expression levels; functional characterization and immunolocalization of glucose and ammonium transporters; NanoSIMS analysis to localize the sites at which new carbon and new nitrogen are incorporated into the holobiont; and experiments in which exogenous glucose was provided to aposymbiotic animals to mimic the supply of glucose from the algae in symbiotic animals. It should be noted that beyond their use here in helping to answer the questions noted above, the extensive datasets on tissue-specific and cell-type-specific gene expression should also be a valuable resource for a variety of future studies. In this regard, however, we do also note the caveat that the (mostly) tentacle-derived data on tissue-specific expression may not in all cases reflect the situation in the rest of the animal.

Previous experiments using rapid tracking of the fate of ^13^C-bicarbonate supplied to symbiotic animals had indicated that newly fixed carbon is transferred from the algae to the host gastrodermal cells primarily as glucose (*29*). Consistent with this hypothesis, we have now shown that the expression of at least six glucose transporters is upregulated in symbiotic relative to aposymbiotic gastrodermal cells. Moreover, the localizations of three such transporters examined are consistent with their putative roles in the transport of glucose across the symbiosome membrane into the cytoplasm of the gastrodermal cells, across the plasma membranes of the gastrodermal cells into the adjacent tissue, or both. Although the resolution of the immunofluorescence images is not sufficient to discriminate among these possibilities, the hypothesis that glucose is trafficked from the gastrodermal cells to the epidermal (and other) cells is also supported by the observations (i) that at least two of the glucose transporters are also significantly upregulated in symbiotic relative to aposymbiotic epidermal cells and (ii) that at least one of the glucose transporters (AipGLUT8α) appears to be localized, in part, to the gastrodermis-epidermis boundary.

Similarly, our analysis of the expression and localization of ammonium transporters suggests that in animals with few or no algal symbionts, the excess ammonium generated by heterotrophic metabolism (*8, 13, 16, 31*) is excreted to the environment at least in part through the bidirectional transporter AipRhBG1 in the epidermal-cell outer membranes. In contrast, when the supply of algal-derived glucose allows abundant incorporation of ammonium into organic compounds, thus reducing intracellular ammonium concentrations (*32*), both the unidirectional transporter AipAMT1 and a substantial fraction of the AipRhBG1 are found at the outer surface of the epidermis, and thus in position to take up environmental ammonium. In addition, another substantial fraction of the highly upregulated AipRhBG1 is found around the gastrodermal cells and at the epidermis-gastrodermis boundary, as also observed in the coral *Acropora yongei* (*33*), and thus in position to distribute both retained and acquired ammonium from the epidermis to the gastrodermal cells and their resident algae.

Despite longstanding appreciation of the importance of conservation and/or recycling of nitrogen for symbiotic cnidarians (*8, 11, 13, 15, 16, 31, 34*), it has remained unclear which element(s) of the holobiont are responsible for these activities. Our results suggest that not only the algae but also both major host tissues are involved. First, our transporter studies suggest that algal-derived glucose and environmental ammonium are both available to both the gastrodermal and epidermal cells of the host as well as to the algae. Second, in agreement with earlier studies of GS enzymatic activities (*8*) and of GS and GOGAT mRNA levels (*14–16*), we found both mRNAs to be highly upregulated at the whole-organism level in symbiotic animals, while no significant difference was found between gastrodermis and epidermis in this regard, suggesting that both major host tissues participate in ammonium incorporation by the GS/GOGAT system during symbiosis. Finally, the NanoSIMS data show a robust and highly coordinated incorporation of both ^13^C and ^15^N in both major tissues of the host, with a total incorporation on a par with that in the algae. These findings are consistent with a recent metabolomic study showing that carbon and nitrogen are significantly co-integrated into amino acids by different symbiotic cnidarians, including Aiptasia and the coral *Stylophora pistillata* (*32*). Moreover, the observation that the total ^15^N incorporation was actually greater in the epidermis than in the gastrodermis argues against the possibility that the organic-nitrogen compounds are all synthesized in the algae and then passed to the host.

It had seemed possible that the provision of glucose by the algae to the host was all that was needed to trigger the entire suite of changes in gene expression, metabolic function, and cellular organization that distinguish a symbiotic anemone from an aposymbiotic one. Indeed, when we provided exogenous glucose to aposymbiotic animals, some changes in gene expression (and presumably in the metabolic pathways governed by those gene products) mimicked those in symbiotic animals. Most notably, the upregulation of GS and GOGAT was essentially identical in the two cases, indicating that the metabolic response to incorporate more ammonium needs only an abundant supply of glucose to be triggered. However, many other changes in gene expression that are seen in symbiotic animals (notably the upregulation of the AipRhBG1 ammonium transporter), as well as the relocalization of ammonium transporters, were not reproduced in the aposymbiotic animals provided with exogenous glucose. Thus, the algae must provide at least one other signal to the host that promotes the additional responses needed for an effective symbiosis. Disruption of such a signal(s) under stress conditions might affect the expression (*35*) and/or localization of symbiosis-associated nutrient transporters and thus disrupt the coordinated incorporation of carbon and nitrogen (*36*), leading to the breakdown of nutrient cycling and symbiosis. Hence, it will clearly be of great interest to determine the nature of this signal(s).

In summary, we have used a combination of genomic, genetic, cell-biological, physiological, and biophysical methods to clarify several previously obscure but very important aspects of the interaction between the host and the algae in the cnidarian-dinoflagellate symbiosis. Most notably, our data indicate that both major host tissues and the symbiotic algae all participate in the critical conservation and recycling of nitrogen, a resource that is typically limiting for growth in the coral-reef environment.

## Author contributions

M.A. and G.C. conceived the study, coordinated with all the coauthors, and supervised the whole project. G.C. performed the tissue-specific RNA-seq experiments. G.C., M.A., J.S.P., P.A.C. Y.J.L, C.J.K., V.S., C.R.V., and V.M.W. interpreted the tissue-specific data. G.C., L.E., and H.Z. performed single-cell RNA-seq and analyzed the data. G.C., M.K.K., L.L., B.H., J.M., P.A.C., C.J.K., H.H., J.R.P, and M.A. performed immunofluorescence experiments. G.C. and O.R.S. conducted the yeast experiments. J.-B. R., N.R., G.C., M.J.C., P.G., J.B., and M.P. performed the NanoSIMS experiments and analyzed the data. G.C. and M.A. wrote the initial draft with inputs from all authors. G.C., M.A., and J.R.P finalized the manuscript and integrated all other author’s comments. All authors reviewed the manuscript.

## Data availability

All data needed to evaluate the conclusions are present in the paper and/or the Supplementary Materials. Sequencing data were deposited in the NCBI Sequence Read Archive under BioPoject codes: PRJNA631577, PRJNA879255, and PRJAN879277.

## Supplementary Materials

Materials and Methods

References (37–54)

Figs. S1 to S8

Tables S1 to S19

Data S1 to S5

