## Supplementle for "Molecular insights into the Darwin paradox of coral reefs from the sea anemone Aiptasia"

### Materials and Methods

#### Animals and maintenance

Throughout this study, "seawater" refers to autoclaved, filtered seawater from the Red Sea. All experiments in this study used individuals of the sea anemone Aiptasia (sensu *Aiptasia pallida*, *Exaiptasia pallida*, or *Exaiptasia diaphana*) clonal strain CC7 (*37*) maintained in seawater. To generate anemones free of algal endosymbionts, animals were treated by cold-shock at 4 °C for 4 h, followed by ~30 d of treatment in 50 μM Diuron (Sigma-Aldrich) with daily water changes (*15*). At the end of the treatment, all anemones were individually inspected by fluorescence microscopy to confirm the absence of algal chlorophyll. A subset of these aposymbiotic anemones was then inoculated with the symbiotic *Breviolum minutum* algal strain SSB01 (*38*) and cultured until algal numbers were stable. Both symbiotic and aposymbiotic anemones were kept on a 12 h:12 h light:dark cycle with ~40 μmol photons m^-2^s^-1^ of photosynthetically active radiation and fed with freshly hatched brine shrimp (*Artemia salina*) approximately three times per week with seawater changes the day after feeding.

Eight small tanks (~500 ml) were set up for this study, with four tanks for symbiotic and four for aposymbiotic anemones. All tanks contained anemones of similar size (pedal disc ~0.5 cm in diameter) and were maintained in the same incubator for the entire duration of the study. Prior to any sampling, the animals were not fed for 3 d to prevent contamination with genetic material from the brine shrimp. Aposymbiotic anemones were confirmed to be free of algae by fluorescence microscopy immediately before sampling for experiments.

#### Laser microdissection

One anemone from each of the eight 500-ml tanks was collected at 11:00 AM (~6 h into the light period), immediately snap-frozen in liquid nitrogen, embedded in Tissue Freezing Medium (Electron Microscopy Sciences), and stored at -80 °C until cryosectioning. The cryostat (CM3050 S, Leica) was pre-chilled to a chamber temperature of -23 ℃, and samples were equilibrated to the chamber temperature for 20 min, then sectioned at a thickness of 8 μm. Sections were then transferred onto pre-cooled RNase-free polyester (POL) membrane metal frame slides (Leica) and air-dried for 5 min in a RNase-free fume hood.

Gastrodermal (Gas) and epidermal (Epi) cell layers were identified in sections (mostly from tentacles) at both 10× and 20× magnifications using a Leica LMD 6000 microscope with Leica filter cubes B/G/R and A. Each region of interest was traced individually using the laser-microdissection (LMD) software and dissected using the ultraviolet laser beam. The dissected pieces were collected in CapSure Macro LCM Caps (Thermo Fisher) containing 40 μl RNA-extraction buffer from the Arcturus PicoPure RNA Isolation Kit (Thermo Fisher). The harvested cells were lysed by incubation at 42 °C for 30 min, vortexed briefly, then kept at -80 °C until further processing.

**Glucose supplementation**

To test the effect of exogenous glucose on gene expression, we compared the transcriptomic profiles of glucose-supplemented aposymbiotic anemones with those of symbiotic and aposymbiotic animals without glucose supplementation. Experiments were performed using three wells in each of five 6-well plates; each well contained 8 ml of seawater. Each plate contained one symbiotic and two aposymbiotic anemones in separate wells, and glucose was added to one of the wells containing an aposymbiotic anemone at a final concentration of 10 mM. Seawater was changed every 2 d, and a new glucose dose was added after each water change. To avoid batch effects, the five plates, representing five biological replicates, were processed simultaneously. The whole experiment lasted for 11 d with anemones being collected on the last day at 11:00 AM, ~6 h into the light period. The collected animals were snap frozen in liquid nitrogen and kept at -80 °C until further processing for whole-animal RNA-Seq analysis.

#### Isolated-tissue and whole-animal RNA-Seq

For the LMD-isolated tissue samples, total RNA from each cell lysate was extracted using the Arcturus PicoPure RNA Isolation Kit following the protocol for use with CapSure Macro LCM Caps. The quality of each RNA sample was assessed on an Agilent 2100 Bioanalyzer using the Agilent RNA 6000 Pico Kit. cDNA was synthesized using the Ovation RNA-Seq System V2 kit (NuGen) following the manufacturer’s instructions. The amplified cDNA was fragmented to ~200 bp by shearing using a Covaris ultrasonicator according to the manufacturer's suggested protocol. The fragmented cDNA was used to generate multiplexed sequencing libraries (mean insert size 200 to 250 bp) using the NEBNext Ultra II DNA Library Prep Kit for Illumina sequencing. The samples were pooled and sequenced on four lanes of an Illumina HiSeq 2000 to generate paired-end reads.

For samples collected from the glucose-supplementation experiment, total RNA was extracted using the RNeasy Mini Kit (Qiagen) following the manufacturer’s protocol for extraction from animal tissues. RNA quality was then assessed as described above. RNA-Seq libraries were prepared using the TruSeq RNA Library Preparation Kit v2 (Illumina) according to the manufacturer’s instructions. All 15 libraries were then pooled and sequenced on a S1 flow cell with an Illumina Novaseq 6000.

RNA-Seq reads were mapped against the revised Aiptasia gene models (*39*) to quantify their expression levels using kallisto v0.44.0 (*40*). For the LMD-isolated tissue samples, analysis of differential gene expression was performed using sleuth v0.29.0 (*41*) with two schemes. First, four pairwise comparisons (SymGas *vs*. ApoGas; SymEpi *vs*. ApoEpi; SymGas *vs*. SymEpi; and ApoGas *vs*. ApoEpi) were performed to identify the main drivers of gene-expression changes. Second, multiple-factor analysis including both tissue source and symbiotic state was applied to determine their effects on the expression profiles. The differentially expressed genes (DEGs) identified in the latter scheme were further clustered into five modules based on hierarchical clustering. Gene Ontology (GO)-term enrichment analysis was conducted on the genes in these modules using topGO (*42*) as described previously (*15*).

In the glucose-supplementation experiments, differential-expression analysis was done only with the pairwise scheme. Expression levels of DEGs identified from each pairwise comparison were expressed as transcripts per million (TPM). The gene-expression levels in each condition were compared and visualized using a ternary plot (*43*).

#### Single-cell RNA-Seq, data clustering, and marker identification

Single-cell RNA-Seq was performed to determine cell-type-specific gene-expression profiles for all Aiptasia cell types. One symbiotic and one aposymbiotic anemone were rinsed with 10 ml of phosphate-buffered saline (PBS, Merck) and dissected in 5 ml of the same buffer containing 100 μg/ml Liberase (Roche, SKU 5401119001) at 23 °C for 1 h. Each cell suspension was filtered through a 40-µm Falcon cell strainer (Thermo Fisher), adjusted to a density of ~2,000 cells/μl, and run directly on the 10× Chromium system (10× Genomics). Single-cell libraries were generated using the 10× Chromium Single Cell 3' Reagent Kit V3 according to the manufacturer’s instructions. Library size distribution and concentration were determined using the Agilent Bioanalyzer with the High Sensitivity DNA kit and the QuantStudio 3 real-time PCR system (Thermo Fisher) with the KAPA DNA-quantification kit (Roche), respectively. The libraries were sequenced using an Illumina HiSeq 4000 to generate paired-end reads.

Reads from the single-cell RNA-Seq were processed using kallisto v0.46.2 (*40*) and BUStools v0.40.0 (*44*) with a wrapper for single-cell data (*45*). The resulting expression matrices were then read into R and analyzed subsequently using Seurat v3.2.0 (*46*). Cells with <200 unique molecular identifiers (UMIs) were removed from further analyses.

The data derived from different samples were integrated using Seurat with the batch-effect-removal workflow. Briefly, the 2,000 genes with the highest dispersion in each sample were identified and used as anchor features for data integration. The first 30 dimensions were included as a neighbor-search space to find connections between samples. The cell clusters were classified at a resolution of 0.9 following the Leiden algorithm (*47*). The marker genes for each cell cluster were identified using a Wilcoxon Rank Sum test (adjusted *p* value < 0.05), the default test specified in the *FindAllMarks* function implemented in Seurat. The expression patterns of these marker genes were visualized with either Seurat (*46*) or Scanpy (*48*).

#### Cell-cluster annotation and tissue-data deconvolution

To integrate the single-cell and tissue-specific RNA-Seq data, we performed a deconvolution analysis on the tissue-specific data using MuSiC (*21*). This method uses the gene expression information from single-cell RNA-Seq data to estimate the cell composition of tissues based on their transcriptomic profiles.

To define marker genes for cell-cluster identification in Aiptasia, we used previously identified markers from the soft coral Xenia (*23*). Orthologous gene groups between Xenia and Aiptasia were identified using OrthoFinder (*49*). The overlaps between cell markers in these two species were examined in R to assist with Aiptasia cell-cluster annotation. To better understand the dominant function of each cell cluster, we performed a gene-set-variation analysis (GSVA) to determine the activities of annotated pathways in Aiptasia following a previously described workflow (*50*). The activities of the top five enriched pathways represented by GSVA scores were then visualized using ComplexHeatmap (*51*).

#### Generating and verifying antibodies against glucose and ammonium transporters

To examine the tissue and cellular localizations of the major Aiptasia transporters for glucose (AipSLC2A8α, AIPGENE2706; AipGLUT1α, AIPGENE12082; AipGLUT8α, AIPGENE18406) and ammonium (AipAMT1, AIPGENE17420; AipRhBG1, AIPGENE18105), we generated rabbit polyclonal (AipSLC2A8α, AipGLUT1α. AipAMT1, and AipRhBG1; GenScript) or mouse monoclonal (AipGLUT8α; Abmart) antibodies against antigen peptides that are specific to each of the proteins (Table S18).

To verify the specificities of the antibodies by Western blotting (Fig. S5), Aiptasia cell lysates were prepared from ~20 anemones by homogenizing in ice-cold NP40 cell-lysis buffer (Thermo Fisher, #FNN0021) using TissueLyser II (Qiagen). After determining the protein concentrations of freshly prepared homogenates using a DC protein assay kit (Bio-Rad), 20 μg (for AipSLC2A8α and AipGLUT1α), 15 μg (for AipGLUT8α), or 10 μg (for AipAMT1 and AipRhBG1) of total protein per sample were then resolved on a 10% SDS-PAGE gel and transferred onto a 0.45-μm PVDF membrane using a wet/tank blotting system (Bio-Rad). The membrane was then blocked in 5% non-fat milk for 2 h at ~23 °C while samples of the antibodies were incubated with either TBST (20 mM Tris, 150 mM NaCl, pH 7.6, 0.1% Tween 20) or TBST supplemented with the appropriate antigen peptide (400 μM) for 2 h at 4 °C. The blocked membrane was then treated with the pre-incubated antibody (1:1,000 in 5% non-fat milk) at 4 °C overnight and then incubated with HRP-conjugated goat anti-mouse-IgG (1:20,000; for anti-AipGLUT8α) or HRP-conjugated goat anti-rabbit-IgG secondary antibody (1:5,000; for the other primary antibodies) for 1 h at ~23 °C. The immunoblots were then visualized by chemiluminescence using the ECL Western-blotting-substrate kit (Bio-Rad) and a ChemiDoc XRS+ system (Bio-Rad). The Western-blot-verified antibodies were then used for immunofluorescence staining experiments.

#### Immunofluorescence staining of glucose and ammonium transporters

Anemones (symbiotic, aposymbiotic, or aposymbiotic treated with glucose) were relaxed in autoclaved seawater containing 3.75% (w/v) MgCl_2_ for ~15 min, then fixed overnight in freshly prepared 4% paraformaldehyde at 4 °C. The fixed animals were dehydrated with a series of washes: PBS, 5 min; 50% ethanol, 5 min, twice; 70% ethanol, 5 min, twice; 80% ethanol, 5 min, twice; 90% ethanol, 5 min, twice; 100% ethanol, 10 min, twice; and xylene, 15 min, twice. The dehydrated specimens were then embedded in paraffin and cut into 5- or 10-μm sections (some of tentacles, some of body stalk) using a Leica RM2125 microtome. The sections were collected on glass slides and then dewaxed and rehydrated with seven steps of reversed washes: xylene; 100%, 90%, 80%, 70%, and 50% ethanol; and PBS containing 0.1% Triton X-100 to permeabilize the cells. After blocking at ~23 °C for 30 min using a blocking buffer containing 5% normal goat serum, 1% BSA, and 0.3% Triton X-100 (anti-AipSLC2A8α, anti-AipGLUT1α, anti-AipAMT1, and anti-AipRhBG1) or 5% normal goat serum, 1% BSA, and 0.1% Tween20 (anti-AipGLUT8α), the sections were then incubated overnight at 4 °C with the appropriate primary antibodies (for AipSLC2A8α, AipGLUT1α, AipAMT1, and AipRhBG1: 10 μg/ml in PBS containing 1% normal goat serum and 0.3% Triton X-100; for AipGLUT8α: a 1:500 dilution in the blocking buffer described above). Secondary antibodies [for AipSLC2A8α, AipGLUT1α, AipAMT1, and AipRhBG1: Alexa Fluor 555-conjugated goat anti-rabbit-IgG (ab150078, Abcam) at 4 μg/ml in PBS containing 1% normal goat serum and 0.3% Triton X-100; for AipGLUT8α: a 1:1000 dilution of Alexa Fluor 488-conjugated goat anti-mouse-IgG (ab150113, Abcam) in the blocking buffer described above] were then applied to the sections for 1 h at ~23 °C. The slides were washed briefly three times with PBS, stained for 2 min with 5 μg/ml Hoechst 33342 (H3570, Thermo Fisher), mounted with ProLong Diamond Antifade Mountant (P36961, Thermo Fisher), and sealed with clear nail polish.

Fluorescence images showing tissue or cellular localizations of the transporters were then captured using a Leica Stellaris 8 FALCON (anti-AipSLC2A8α and anti-AipGLUT1α), a Keyence BZ-810 microscope (anti-AipGLUT8α), or a Leica TCS SP8 STED X (anti-AipAMT1 and anti-AipRhBG1). All images were then analyzed using Fiji (*52*).

#### Rescue of yeast mutants with transformed Aiptasia transporters

To test the functions of symbiosis-induced putative glucose and ammonium transporters, we performed yeast-mutant-rescue experiments. For the glucose transporters, we amplified full-length transcripts for AipSLC2A8α, AipGLUT1α, and AipGLUT8α using gene-specific primers (Table S19) and cloned the resulting cDNAs into the yeast vector pSH100 [YCplac33 MET25pro MCP-mCherry (*53*), a gift from Robert Singer & Daniel Zenklusen (Addgene plasmid # 45930; http://n2t.net/addgene:45930; RRID:Addgene_45930)]. The constructs were then transformed into CFY07, a yeast mutant strain that has completely lost hexose-uptake ability due to the concurrent knockout of 20 sugar-transporter genes plus an extracellular trehalase gene (28). For the ammonium transporters, we amplified full-length transcripts for AipAMT1 and AipRhGB1 using gene-specific primers (Table S19) and cloned the resulting cDNAs into the yeast vector pYES2.1/V5-His-TOPO (K415001, Thermo Fisher). The resulting plasmids were sequenced to confirm sequence fidelity and transformed into the ammonium-transporter-deficient mutant yeast strain 31019b, which lacks the three yeast ammonium transporter genes (*MEP1*, *MEP2*, and *MEP3*), at least one of which is essential for yeast to grow using ammonium as sole nitrogen source (*27*).

In all cases, yeast strains were grown overnight to exponential phase in complete medium [YPM (1% yeast extract, 2% peptone, 2% maltose) for CFY07 and YPD (1% yeast extract, 2% peptone, 2% glucose) for 31019b], pelleted from 1 ml of each culture, washed with MilliQ water, and resuspended to a concentration of 1 × 10^4^ cells/μl. To test growth, dilution series were then prepared and plated on the appropriate synthetic medium, resulting in five colonies for each strain containing approximately 50000, 5000, 500, 50, and 5 cells initially. CFY07 and its transformants were plated on YNB medium containing 2% glucose, whereas 31019b and its transformants were plated on Difco Yeast Nitrogen Base medium without amino acids and ammonium (YNB-N) containing 20 mM NH_4_Cl and 2% galactose. Images showing cell growth were acquired after incubation at 30 ℃ for 2 d.

To characterize the directionality of the ammonium transporters, the same 31019b transformants were plated on YNB-N medium containing 2% galactose, 200 mM methylammonium, and 0.1% proline. Proline can serve as the nitrogen source to support cell growth, but methylammonium is a toxic ammonium analogue. Hence, cells expressing a unidirectional ammonium transporter accumulate the lethal chemical over time and die, whereas a bidirectional transporter can remove toxic methylammonium from the cells and thus allow them to survive. Images showing cell growth on the methylammonium plates were acquired 3 d after inoculation.

#### Isotope labeling and NanoSIMS analysis

To investigate the dynamics of carbon and nitrogen assimilation in symbiotic and aposymbiotic animals, we performed stable-isotope-labeling experiments on two anemones of each type. Individual animals were incubated for 24 h in 25-ml incubation chambers containing artificial seawater (ASW) at 25 °C under a 12:12 h light-dark cycle with a light intensity of 40 μmol photons m^-2^ s^-1^. The ASW was freshly prepared and contained 20.8 g/L NaCl, 4.4 g/L MgCl_2_, 3.5 g/L Na_2_SO_4_, 1 g/L CaCl_2_, 0.59 g/L KCl at pH 8.2. It was supplemented with NaH^13^CO_3_ (> 98 atom % ^13^C) and ^15^NH_4_Cl (>98 atom % ^15^N) at final concentrations of 2 mM and 5 μM, respectively. Following the 24-h incubations, all specimens were immediately transferred to a fixative solution (1.25% glutaraldehyde plus 0.5% paraformaldehyde in 0.1 M sodium phosphate buffer, pH 7.5) and stored at 4 °C for 24 h before further processing.

Several individual tentacles were then collected from each animal using a stereomicroscope, post-fixed for 1 h at ~23 °C in 1% OsO_4_ in 0.1 M sodium phosphate buffer (pH 7.5), and dehydrated using a series of rinses with increasing ethanol concentrations (50, 70, 90, and 100%) followed by 100% acetone. Tissues were then gradually infiltrated using Spurr’s resin (Electron Microscopy Sciences) at increasing concentrations (25, 50, and 75% in ethanol; then 100%) and finally embedded in 100% resin. Semi-thin sections (150 nm) were cut using a Leica Ultracut E microtome, mounted on silicon wafers (ProsciTech), and gold-coated using a Quorum Q150T sputter coater.

The gold-coated sections were then imaged using the NanoSIMS 50 ion probe at the Centre for Microscopy, Characterisation, and Analysis at the University of Western Australia. The surfaces of the samples were bombarded with a 16 keV primary Cs^+^ beam focused to a spot size of about 100 nm, with a current of ∼2 pA. Secondary molecular ions ^12^C^12^C^−^, ^12^C^13^C^−^, ^12^C^14^N^−^, and ^12^C^15^N^−^ were collected simultaneously in electron multipliers at a mass resolution (M/ΔM) > 8,000, which is sufficient to resolve isobaric species (e.g., ^12^C^13^C^−^ from ^12^C^12^C ^1^H^−^ and ^13^C^14^N^−^ from ^12^C^15^N^−^). Large mosaics of the different cell layers comprising, on average, 16 ± 3 images (45 μm raster with 512 × 512 pixels) were recorded for all targeted secondary ions by rastering the primary beam across the sample with a dwell-time of 10–20 ms per pixel. After drift correction, the ^13^C/^12^C and ^15^N/^14^N maps were generated based on ^12^C^13^C^−^/^12^C^12^C^−^ and ^12^C^15^N^−^/^12^C^14^N^−^ ratios as hue-saturation-intensity images (HSI), where the color scale represents the isotope ratio, and further analyzed using Fiji with the Open-MIMS plug-in (<https://github.com/BWHCNI/OpenMIMS/wiki>). Isotopic enrichments in the Aiptasia mosaics were quantified in the epidermis, gastrodermis (excluding the algal cells in symbiotic samples; Fig. S8), and algal cells by extracting data from each pixel (spatial resolution ~88 nm) in the regions of interest across the tentacle cross sections. Over 10^6^ pixel values were extracted for each tissue layer. To eliminate potential effects on the subsequent statistical analyses that might be caused by different sample sizes (total pixel numbers), the pixel values from each tissue layer were randomly assigned into 10,000 bins. The ^13^C/^12^C and ^15^N/^14^N ratios were then used to quantify the isotope enrichment for each bin. The Pearson correlations between ^13^C and ^15^N enrichments were calculated based on the relative signal intensities for each treatment and tissue layer, and the isotope-enrichment levels from different tissues were compared using R package *cocor* (*54*). In addition, the same NanoSIMS mosaics were used to approximate the relative overall contributions by the anemone tissues and the algae to ammonium assimilation in the holobiont. For this, the total ^15^N counts across the regions of interest were corrected for the background levels of natural-isotope abundance (based on isotope ratios from yeast and unlabeled Aiptasia samples) to quantify the excess ^15^N assimilation by each symbiotic partner.

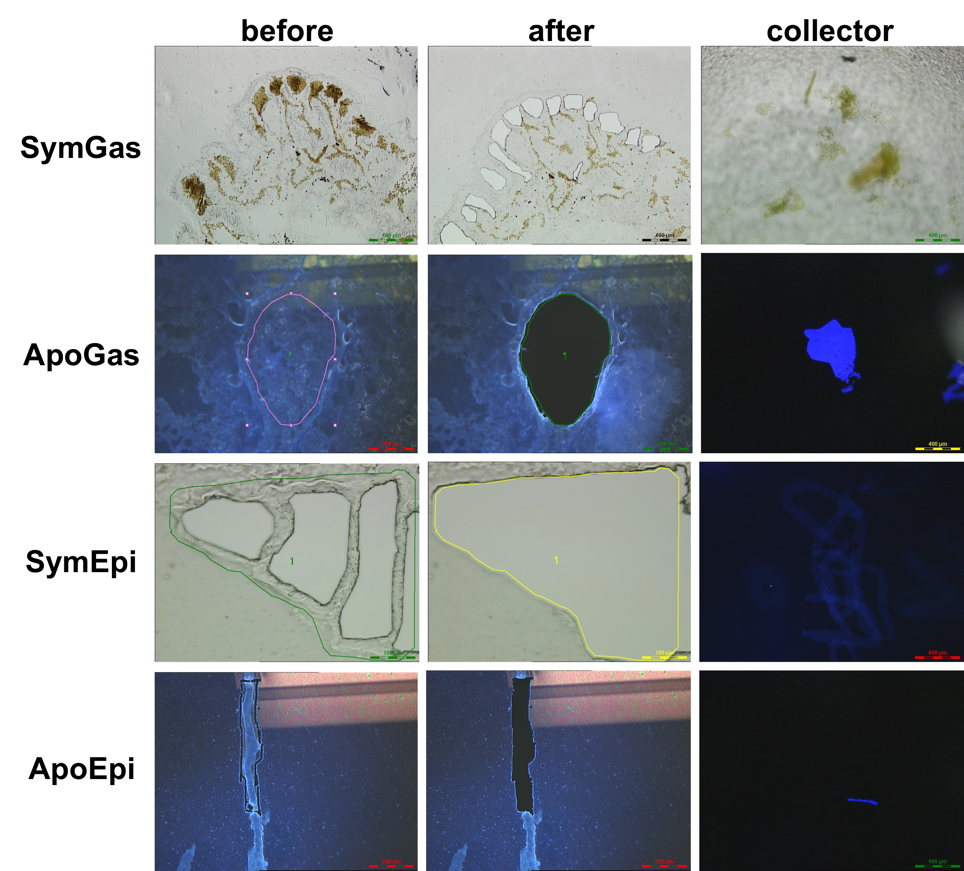

**Fig. S1.** Laser microdissection (LMD) of symbiotic and aposymbiotic Aiptasia. Symbiotic samples were imaged under white light, whereas aposymbiotic samples were visualized using a Leica filter cube B/G/R or A, which helped in identifying their epidermal cell layer. Sym, symbiotic; Apo, aposymbiotic; Gas, gastrodermis; Epi, epidermis.

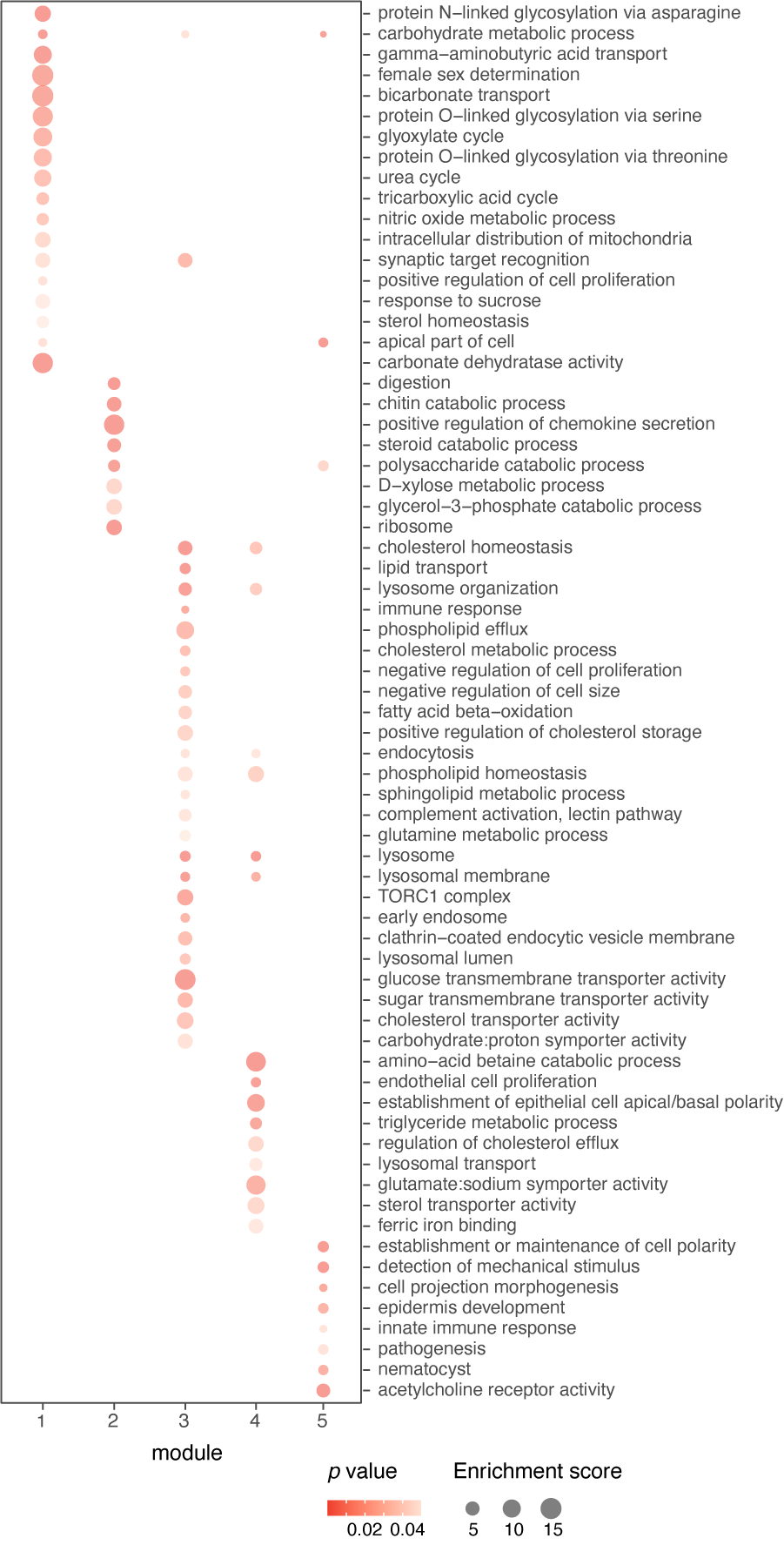

**Fig. S2.** Gene-ontology (GO) terms enriched by the genes in each of the five modules identified from multi-factorial differential-expression analysis (Fig. 1C). The size of each dot represents the enrichment score that was calculated by dividing the actual by the expected number of DEGs associated with the corresponding term.

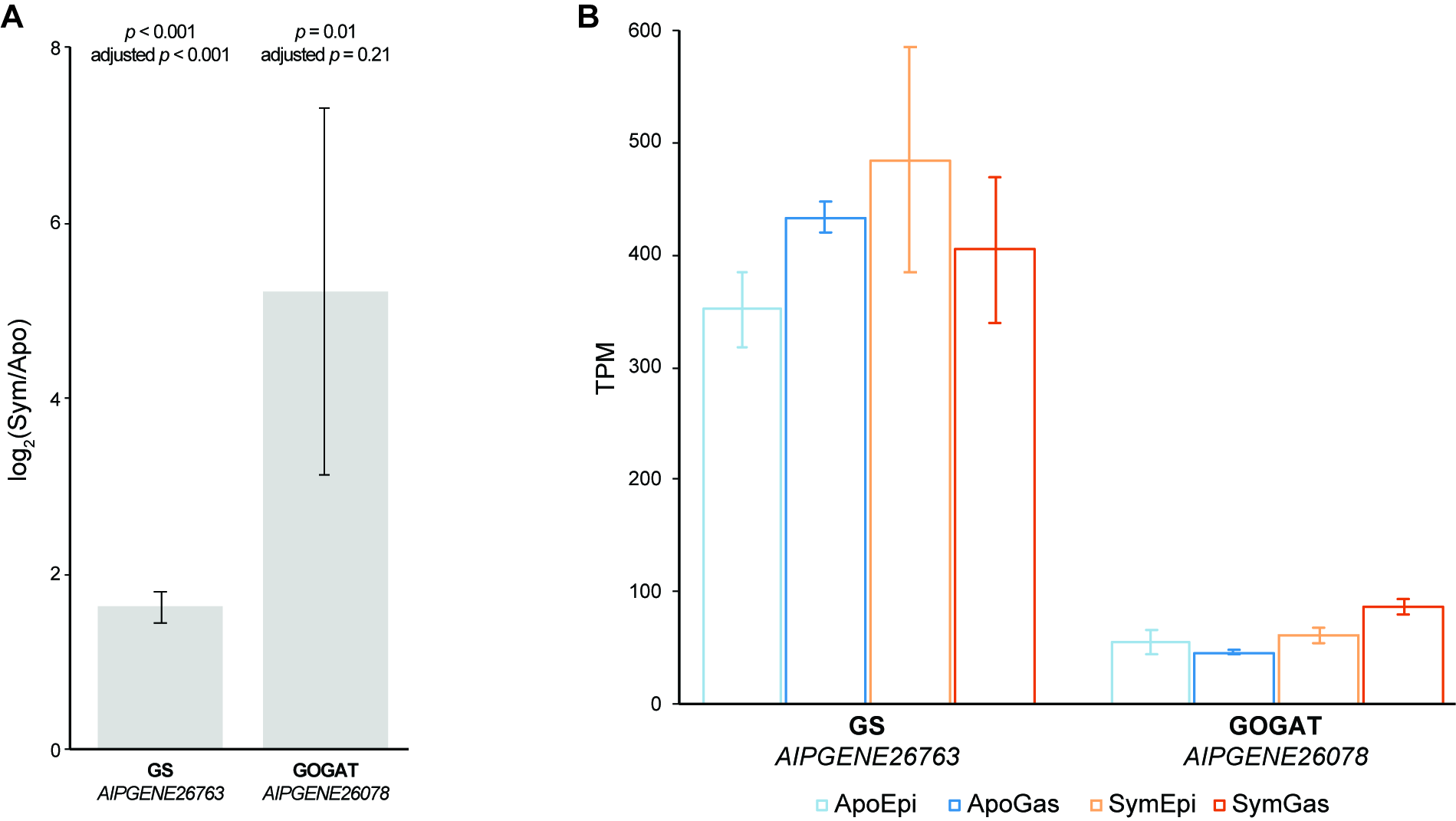

**Fig. S3.** Expression patterns of key genes associated with ammonium assimilation at organism (A) and tissue (B) levels. Organism-level data in (A) were extracted from Cui et al., 2019 (<https://doi.org/10.1371/journal.pgen.1008189.s005>). GS, glutamine synthetase; GOGAT, glutamate synthase. Error bars represent mean ± SE.

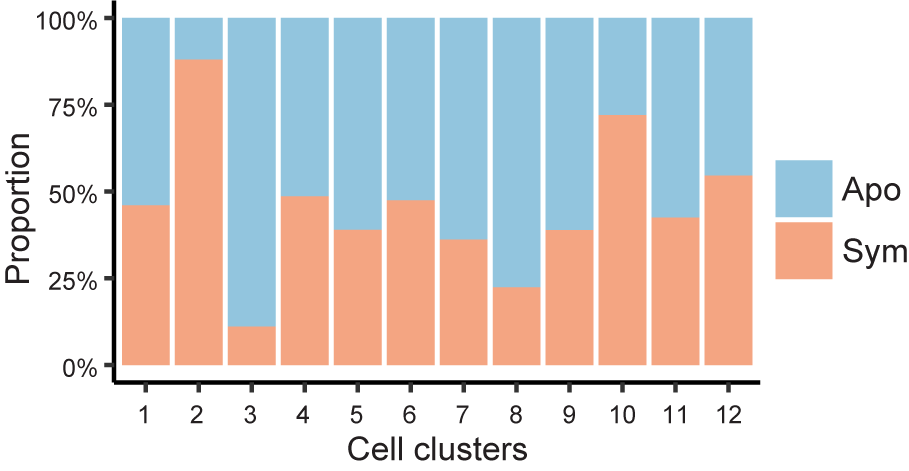

**Fig. S4.** Proportions of cells originating from aposymbiotic or symbiotic anemones in the 12 clusters identified by single-cell RNA-Seq (see **Fig 1, D and E**). Apo, aposymbiotic; Sym, symbiotic.

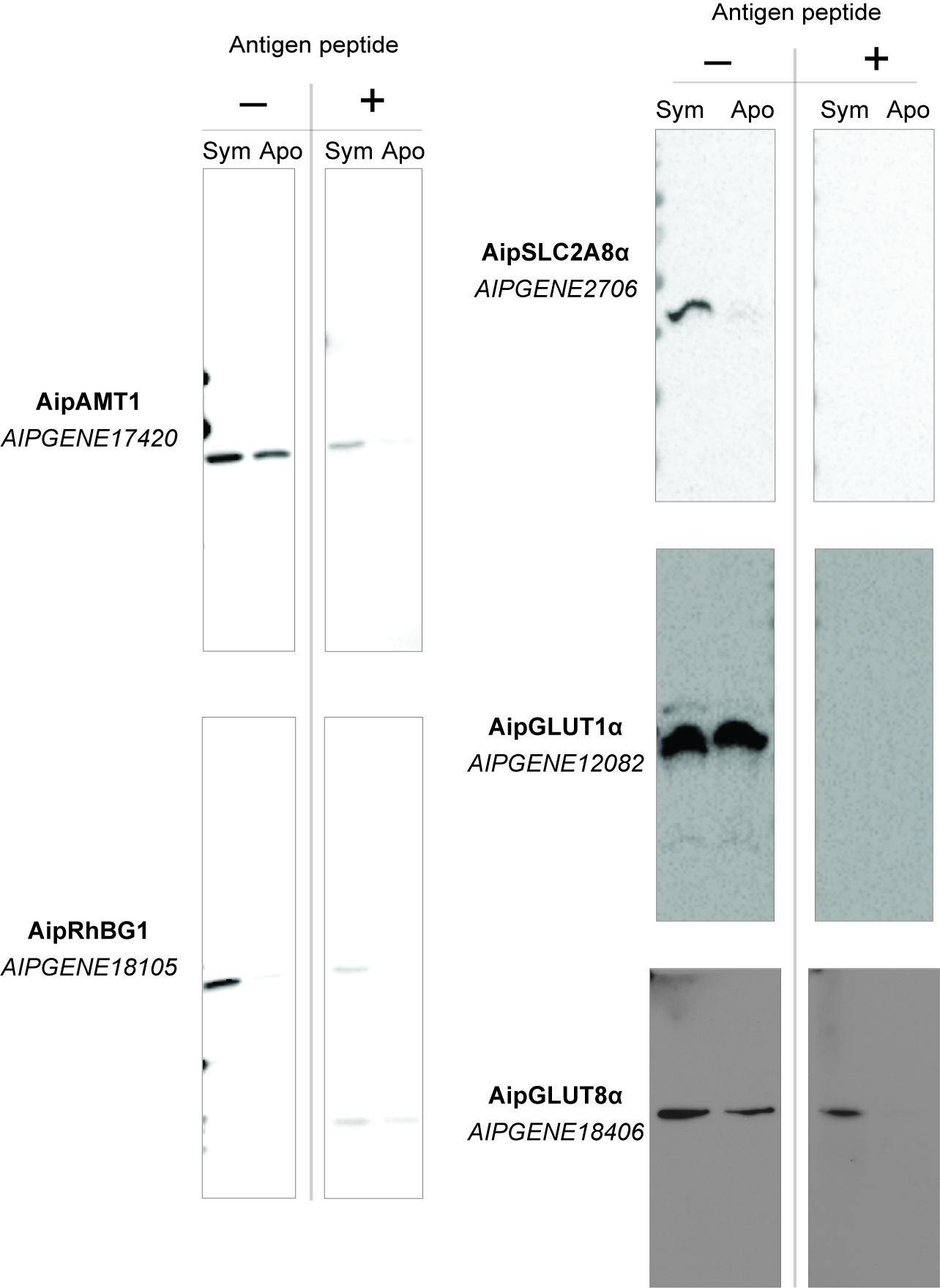

**Fig. S5.** Western-blot validation of the custom antibodies. 10 μg (for AipAMT1 and AipRhBG1), 15 μg (for AipGLUT8α), or 20 μg (for AipSLC2A8α and AipGLUT1α) of total protein isolated from symbiotic (Sym) or aposymbiotic (Apo) anemones were resolved on SDS-PAGE gels and transferred onto PVDF membranes. The membranes were incubated either with only the appropriate antibody (-) or with the appropriate antibody pre-absorbed with its antigen peptide (+). Further details are in Materials and Methods.

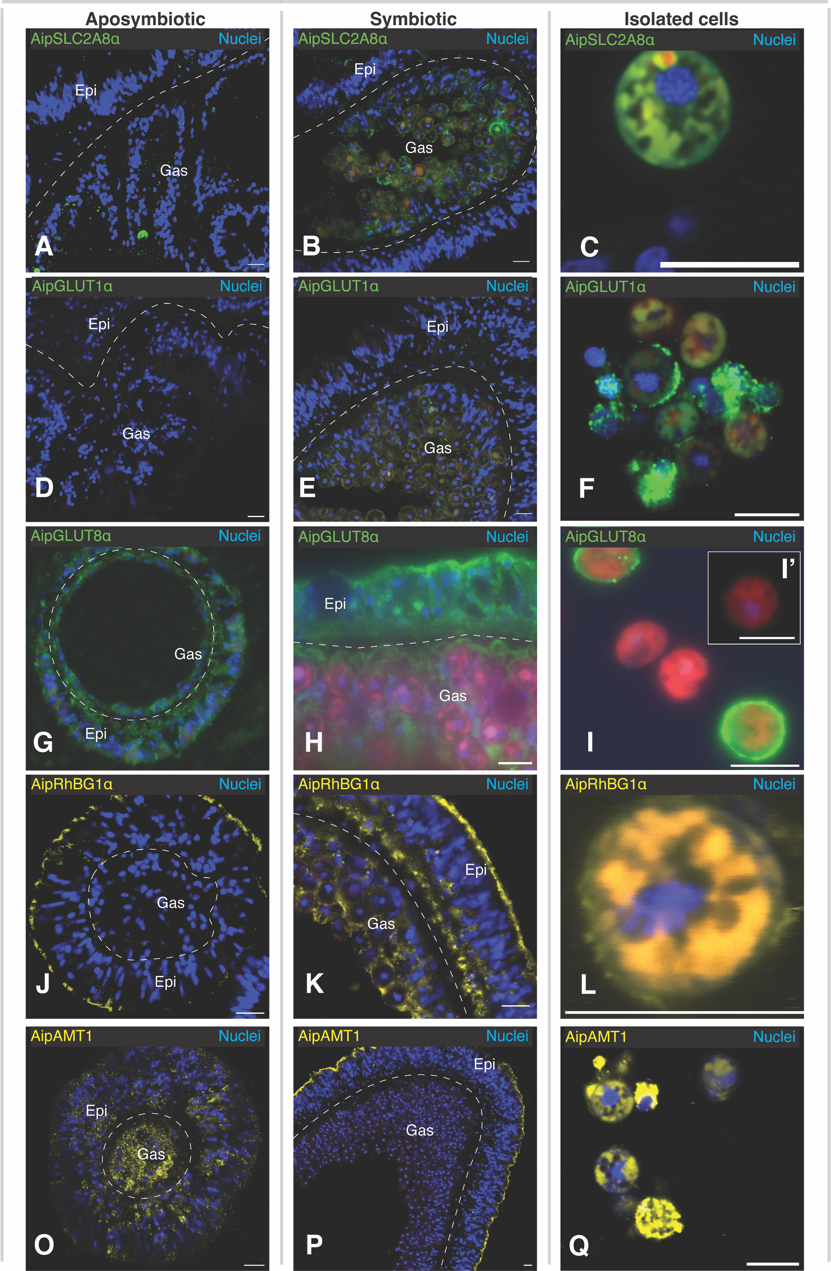

**Fig. S6.** Additional immunofluorescence images (as in **Fig. 2**, E-S) showing relocalization of glucose (A-I) and ammonium (J-Q) transporters induced by symbiosis. Tissue sections of aposymbiotic (A, D, G, J, O) and symbiotic (B, E, H, K, P) anemones are shown along with cells isolated from symbiotic animals (C, F, I, L, Q). Immunofluorescence staining of cultured algal cells (strain SSB01) with the anti-AipGLUT8α antibodies is also shown (I’); the image shown is representative of many such cells observed. Scale bars (all panels), 10 μm.

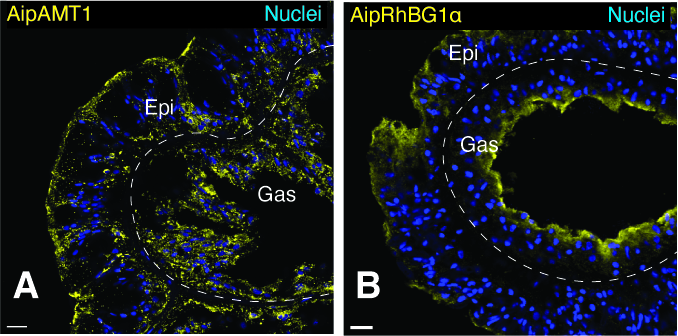

**Fig. S7.** Additional images showing localization patterns of ammonium transporters AipAMT1 (A) and AipRhGB1 (B) in glucose-treated aposymbiotic Aiptasia (see **Fig. 3 C and D**). Scale bars (both panels), 10 μm.

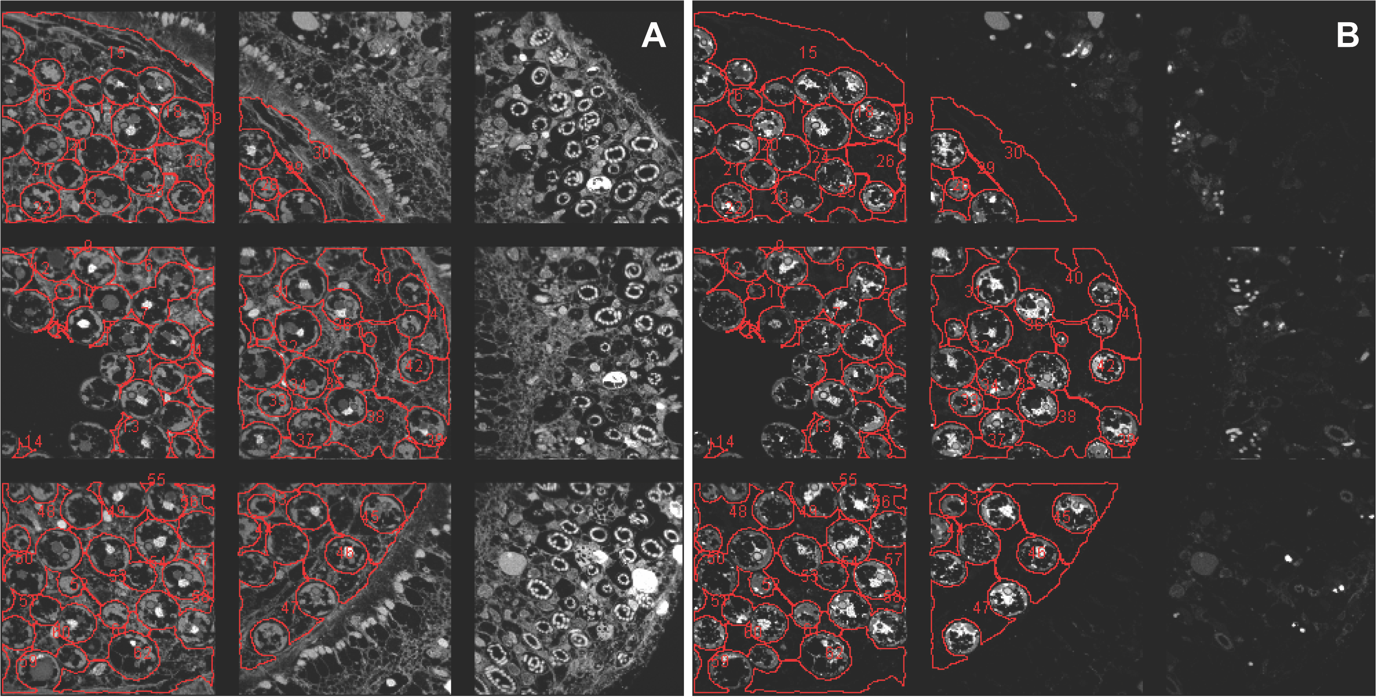

**Fig. S8.** Representative regions of interest (ROI; numbered red outlines) in the gastrodermis displayed for (A) ^14^N and (B) ^15^N. The ROI were drawn manually to include only gastrodermal tissue and not algal-symbiont cells.

**Table S1. Module 1 genes involved in carbon cycling and carbohydrate metabolism**

| **Gene ID**^a,b^ | **Average expression level (TPM)**^c^ | | **Fold change**^c^ | | **Annotation** |
| --- | --- | --- | --- | --- | --- |
|  | **SymEpi** | **SymGas** | **SymEpi/ApoEpi** | **SymGas/ApoGas** |  |
| AIPGENE17854 | 51 | 110 | 2.4 | 6.0 | Isocitrate lyase |
| AIPGENE21331 | 85 | 220 | 1.3 | 5.7 | Carnitine O-palmitoyltransferase 1 |
| AIPGENE15469 | 39 | 53 | 1.5 | 5.4 | Carbonic anhydrase 7 |
| AIPGENE6376 | 59 | 76 | 3.4 | 4.5 | Phosphorylase b kinase gamma catalytic chain PHKG2 |
| AIPGENE2308 | 40 | 56 | 1.6 | 4.2 | Polypeptide N-acetylgalactosaminyltransferase 1 |
| AIPGENE12054 | 26 | 31 | 1.9 | 3.7 | Glucose transporter type 1 |
| AIPGENE12939 | 15 | 26 | 1.6 | 3.7 | Alpha-N-acetylgalactosaminidase NAGA |
| AIPGENE9860 | 50 | 53 | 2.8 | 3.6 | [Pyruvate dehydrogenase (acetyl-transferring)] kinase isozyme 2 PDK2 |
| AIPGENE8506 | 25 | 34 | 2.0 | 3.0 | Solute carrier family 26 member 6 SLC26A6 |
| AIPGENE21162 | 19 | 24 | 1.6 | 2.9 | Carbohydrate sulfotransferase 1 |
| AIPGENE1116 | 19 | 30 | 1.8 | 2.7 | N-acetylglucosamine-6-phosphate deacetylase amdhd2 |
| AIPGENE17843 | 50 | 64 | 2.6 | 2.6 | Malate synthase |
| AIPGENE24851 | 70 | 82 | 1.4 | 2.4 | Serine/threonine-protein kinase/endoribonuclease IRE1A |
| AIPGENE12778 | 49 | 68 | 1.3 | 2.2 | Galactokinase GALK1 |
| AIPGENE5850 | 53 | 50 | 1.1 | 1.9 | Capsule biosynthesis protein CapA |
| AIPGENE6132 | 20 | 22 | 2.1 | 1.8 | Lysosomal alpha-mannosidase |
| AIPGENE3057 | 62 | 69 | 1.8 | 1.7 | Pyruvate carboxylase |
| AIPGENE22466 | 27 | 32 | 1.5 | 1.7 | UDP-glucose:glycoprotein glucosyltransferase 1 |
| AIPGENE18740 | 36 | 41 | 1.6 | 1.6 | Neutral alpha-glucosidase AB |
| AIPGENE25712 | 34 | 18 | 4.1 | 1.4 | Di-N-acetylchitobiase CTBS |
| AIPGENE28157 | 110 | 62 | 2.1 | 1.3 | Carbamoyl-phosphate synthase [ammonia] CPS1 |

^a^ Ordered by SymGas/ApoGas.

^b^ Cyan highlight, a gene encoding a putative glucose transporter that is upregulated in both gastrodermal and epidermal cells of symbiotic animals but not investigated further in this study; yellow highlight, genes encoding glyoxylate-cycle enzymes.

^c^ Numbers are rounded to two significant digits.

**Table S2. Module 1 genes involved in nitrogen cycling and serine/glycine biosynthesis**

| **Gene ID**^a,b^ | **Average expression level (TPM)**^c^ | | **Fold change**^c^ | | **Annotation** |
| --- | --- | --- | --- | --- | --- |
|  | **SymEpi** | **SymGas** | **SymEpi/ApoEpi** | **SymGas/ApoGas** |  |
| AIPGENE18105 | 20 | 410 | 6.8 | 170 | Ammonium transporter Rh type B |
| AIPGENE18835 | 26 | 28 | 2.0 | 3.7 | Glycine N-methyltransferase GNMT |
| AIPGENE22717 | 48 | 72 | 2.4 | 3.1 | Serine-pyruvate aminotransferase AGXT |
| AIPGENE18470 | 27 | 23 | 1.7 | 1.8 | Argininosuccinate synthase |
| AIPGENE28157 | 110 | 62 | 2.1 | 1.3 | Carbamoyl-phosphate synthase [ammonia] CPS1 |

^a^ Ordered by SymGas/ApoGas.

^b^ Cyan highlight, a gene encoding an ammonium transporter (AipRhBG1) that is investigated further in this study.

^c^ Numbers are rounded to two significant digits.

**Table S3. Module 1 genes involved in post-translational modification**

| **Gene ID**^a,b^ | **Average expression level (TPM)**^c^ | | **Fold change**^c^ | | **Annotation** |
| --- | --- | --- | --- | --- | --- |
|  | **SymEpi** | **SymGas** | **SymEpi/ApoEpi** | **SymGas/ApoGas** |  |
| AIPGENE130 | 45 | 100 | 2.4 | 4.6 | Alpha-1,3-Glucosyltransferase ALG8 |
| AIPGENE16395 | 28 | 26 | 1.5 | 3.8 | Low-salt glycan biosynthesis protein Agl12 |
| AIPGENE8406 | 29 | 17 | 2.0 | 2.5 | Polypeptide N-acetylgalactosaminyltransferase 1 GALNT1 |
| AIPGENE26605 | 16 | 16 | 2.6 | 2.3 | GPI ethanolamine phosphate transferase 3 PIGO |
| AIPGENE15461 | 18 | 18 | 1.8 | 1.9 | Dolichyl-phosphate beta-glucosyltransferase ALG5 |
| AIPGENE4856 | 110 | 90 | 3.2 | 1.8 | Protein transport protein Sec24C |
| AIPGENE6178 | 48 | 49 | 1.9 | 1.8 | Dolichyl-diphosphooligosaccharide--protein glycosyltransferase STT3B |
| AIPGENE2497 | 56 | 53 | 1.9 | 1.6 | Protein transport protein Sec23A |
| AIPGENE27679 | 93 | 97 | 1.5 | 1.2 | Protein transport protein Sec24B |
| AIPGENE16 | 60 | 21 | 3.4 | 1.1 | Alpha-1,3-glucosyltransferase ALG6 |

^a^ Ordered by SymGas/ApoGas.

^b^ Genes potentially involved in N-linked glycosylation are highlighted.

^c^ Numbers are rounded to two significant digits.

**Table S4. Module 2 genes involved in food digestion**

| **Gene ID**^a,b^ | **Average expression level (TPM)**^c^ | | **Fold change**^c^ | | **Annotation** |
| --- | --- | --- | --- | --- | --- |
|  | **ApoEpi** | **ApoGas** | **ApoEpi/SymEpi** | **ApoGas/SymGas** |  |
| AIPGENE9704 | 6.8 | 35 | 8.2 | 58 | Chymotrypsinogen B |
| AIPGENE25876 | 290 | 1500 | 32 | 55 | Chymotrypsin B |
| AIPGENE15455 | 98 | 400 | 12 | 51 | Chitinase 3 |
| AIPGENE2028 | 35 | 150 | 8.1 | 48 | Trypsin-3 |
| AIPGENE8235 | 71 | 380 | 15 | 35 | Anionic trypsin |
| AIPGENE23293 | 200 | 740 | 13 | 27 | Acidic mammalian chitinase |
| AIPGENE24288 | 59 | 210 | 2.6 | 21 | Chymotrypsin-like protease CTRL-1 |
| AIPGENE1364 | 77 | 180 | 3.9 | 19 | Meprin A subunit beta MEP1B |
| AIPGENE2734 | 22 | 93 | 11 | 17 | Chymotrypsinogen B |
| AIPGENE28200 | 30 | 85 | 8.1 | 16 | Acidic mammalian chitinase |
| AIPGENE2019 | 10 | 36 | 4.3 | 15 | Trypsin-3 |
| AIPGENE23157 | 9.3 | 27 | 3.3 | 13 | Chymotrypsinogen B |
| AIPGENE28199 | 14 | 22 | 5.2 | 12 | Acidic mammalian chitinase |
| AIPGENE7785 | 24 | 74 | 6.8 | 9.8 | Chitinase 3 |
| AIPGENE17423 | 8.7 | 24 | 3.1 | 9.6 | Maltase-glucoamylase MGAM |
| AIPGENE26805 | 10 | 35 | 3.2 | 9.1 | Chymotrypsinogen B |
| AIPGENE24686 | 17 | 18 | 1.0 | 3.1 | Beta-hexosaminidase HEX |
| AIPGENE6698 | 52 | 63 | 1.7 | 2.9 | Chymotrypsinogen B2 |

^a^ Ordered by ApoGas/SymGas.

^b^ Genes encoding putative chitinases are highlighted; they should allow digestion of a major exoskeletal component of most cnidarian prey animals.

^c^ Numbers are rounded to two significant digits.

**Table S5. Module 3 genes involved in lysosomal/symbiosomal organization**

| **Gene ID**^a,b^ | **Average expression level (TPM)**^c^ | | **Fold change**^c^ | | **Annotation** |
| --- | --- | --- | --- | --- | --- |
|  | **ApoGas** | **SymGas** | **SymGas/SymEpi** | **SymGas/ApoGas** |  |
| AIPGENE18610 | 17 | 840 | 18 | 50 | Sulfated glycoprotein 1 |
| AIPGENE20198 | 1.3 | 42 | 13 | 32 | Sulfotransferase 1C2A |
| AIPGENE762 | 10 | 160 | 8.0 | 15 | Phospholipase B-like 2 |
| AIPGENE8500 | 14 | 110 | 10 | 8.3 | Gamma-glutamyl hydrolase |
| AIPGENE756 | 12 | 90 | 5.3 | 7.5 | Phospholipase B-like 2 |
| AIPGENE8201 | 19 | 120 | 5.2 | 6.6 | Gamma-glutamyl hydrolase |
| AIPGENE9365 | 19 | 120 | 4.3 | 6.4 | Beta-glucuronidase |
| AIPGENE9310 | 78 | 490 | 7.2 | 6.3 | E3 ubiquitin-protein ligase MARCH1 |
| AIPGENE17332 | 42 | 230 | 3.2 | 5.7 | Regulatory-associated protein of mTOR |
| AIPGENE12986 | 14 | 78 | 4.1 | 5.6 | Bcl-2-like protein 1 |
| AIPGENE2358 | 21 | 110 | 8.2 | 5.3 | Deleted in malignant brain tumors 1 protein DMBT1 |
| AIPGENE17384 | 34 | 170 | 4.4 | 4.9 | Regulatory-associated protein of mTOR |
| AIPGENE1615 | 9.3 | 45 | 5.1 | 4.9 | Vacuolar protein sorting-associated protein 4B |
| AIPGENE13031 | 11 | 51 | 3.3 | 4.5 | Alpha-N-acetylgalactosaminidase |
| AIPGENE9338 | 25 | 93 | 3.8 | 3.8 | Beta-glucuronidase |
| AIPGENE23106 | 45 | 170 | 3.0 | 3.7 | Legumain |
| AIPGENE10665 | 33 | 100 | 5.9 | 3.1 | Sulfotransferase 1C2A |
| AIPGENE8377 | 29 | 84 | 2.7 | 2.9 | Sorting nexin-13 |
| AIPGENE11822 | 33 | 89 | 3.3 | 2.7 | AP-5 |
| AIPGENE16722 | 26 | 62 | 2.0 | 2.4 | Lysosomal thioesterase PPT2-A |
| AIPGENE1266 | 25 | 59 | 2.8 | 2.4 | Somatomedin-B and thrombospondin type-1 domain-containing protein |
| AIPGENE14910 | 31 | 66 | 9.1 | 2.1 | Basement membrane-specific heparan sulfate proteoglycan core protein HSPG2 |
| AIPGENE25198 | 75 | 149 | 1.8 | 2.0 | PTB domain-containing engulfment adapter protein 1 |
| AIPGENE13027 | 46 | 87 | 2.3 | 1.9 | Alpha-N-acetylgalactosaminidase |
| AIPGENE23952 | 6 | 113 | 3.2 | 1.8 | Nischarin |
| AIPGENE27709 | 32 | 60 | 1.5 | 1.8 | Lysosome membrane protein 2 SCARB2 |
| AIPGENE5637 | 28 | 50 | 2.5 | 1.8 | Phospholipase B-like 1 |
| AIPGENE1792 | 18 | 31 | 2.1 | 1.7 | Chitinase domain-containing protein 1 |
| AIPGENE17107 | 25 | 41 | 1.7 | 1.6 | Vacuolar protein sorting-associated protein 33A |
| AIPGENE1844 | 43 | 69 | 3.1 | 1.6 | Steryl-sulfatase |
| AIPGENE20604 | 27 | 42 | 1.7 | 1.6 | H(+)/Cl(-) exchange transporter 7 |

^a^ Ordered by SymGas/ApoGas.

^b^ Some genes encoding lysosomal marker proteins are highlighted.

^c^ Numbers are rounded to two significant digits.

**Table S6. Module 3 genes involved in sugar transport**

| **Gene ID**^a,b^ | **Average expression level (TPM)**^c^ | | **Fold change**^c^ | | **Annotation** |
| --- | --- | --- | --- | --- | --- |
|  | **SymEpi** | **SymGas** | **SymGas/SymEpi** | **SymGas/ApoGas** |  |
| AIPGENE18406 | 10 | 70 | 7.8 | 6.9 | Facilitated glucose transporter GLUT8 |
| AIPGENE27196 | 23 | 140 | 9.9 | 6.2 | Solute carrier family 23 member 2 SLC23A2 |
| AIPGENE2706 | 5.0 | 21 | 9.5 | 4.7 | Solute carrier family 2, facilitated glucose transporter member 8 SLC2A8 |
| AIPGENE6170 | 8.0 | 19 | 2.0 | 2.5 | Solute carrier family 2, facilitated glucose transporter member 8 SLC2A8 |
| AIPGENE12097 | 23 | 49 | 1.6 | 2.2 | Solute carrier family 2, facilitated glucose transporter member 1 SLC2A1 |

^a^ Ordered by SymGas/ApoGas.

^b^ Cyan highlight, genes encoding glucose transporters that are investigated further in this study; yellow highlight, genes encoding putative glucose transporters that are upregulated in symbiotic gastrodermal cells but not investigated further in this study.

^c^ Numbers are rounded to two significant digits.

**Table S7. Module 3 genes involved in sterol transport and homeostasis**

| **Gene ID**^a,b^ | **Average expression level (TPM)**^c^ | | **Fold change**^c^ | | **Annotation** |
| --- | --- | --- | --- | --- | --- |
|  | **ApoGas** | **SymGas** | **SymGas/SymEpi** | **SymGas/ApoGas** |  |
| AIPGENE23673 | 3.7 | 70 | 15 | 19 | Lathosterol oxidase |
| AIPGENE2187 | 24 | 390 | 9.9 | 16 | Low-density lipoprotein receptor-related protein 6 |
| AIPGENE3074 | 46 | 460 | 13 | 10 | Protein NPC2 |
| AIPGENE15716 | 45 | 330 | 4.9 | 7.3 | Prolow-density lipoprotein receptor-related protein 1 |
| AIPGENE13786 | 71 | 450 | 4.4 | 6.3 | Lipid storage droplets surface-binding protein 2 |
| AIPGENE12169 | 8.6 | 49 | 4.0 | 5.7 | Low density lipoprotein receptor adapter protein 1 |
| AIPGENE23723 | 850 | 4000 | 8.5 | 4.7 | Apolipophorins |
| AIPGENE20806 | 23 | 60 | 2.2 | 2.6 | Hormone-sensitive lipase |
| AIPGENE7350 | 22 | 44 | 4.3 | 2.0 | Long-chain-fatty-acid--CoA ligase 5 |
| AIPGENE23362 | 190 | 280 | 1.9 | 1.5 | Sterol regulatory element-binding protein 1 |
| AIPGENE23053 | 31 | 34 | 1.4 | 1.1 | Non-specific lipid-transfer protein SCP2 |

^a^ Ordered by SymGas/ApoGas.

^b^ A cholesterol-transport protein previously characterized in Aiptasia and other cnidarians is highlighted.

^c^ Numbers are rounded to two significant digits.

**Table S8. Module 4 genes involved in digestion**

| **Gene ID**^a,b^ | **Average expression level (TPM)**^c^ | | **Fold change**^c^ | | **Annotation** |
| --- | --- | --- | --- | --- | --- |
|  | **ApoGas** | **SymGas** | **ApoGas/ApoEpi** | **SymGas/SymEpi** |  |
| AIPGENE21470 | 190 | 380 | 2.4 | 6.7 | Proactivator polypeptide |
| AIPGENE18552 | 92 | 89 | 2.1 | 6.4 | Acetylcholinesterase |
| AIPGENE27509 | 65 | 99 | 1.8 | 4.6 | Betaine--homocysteine S-methyltransferase 1 |
| AIPGENE26157 | 680 | 350 | 2.8 | 4.1 | Cathepsin L |
| AIPGENE16849 | 170 | 180 | 2.1 | 3.5 | Cathepsin B |
| AIPGENE10977 | 23 | 21 | 2.2 | 3.3 | Betaine--homocysteine S-methyltransferase 1 |
| AIPGENE203 | 20 | 21 | 2.1 | 3.2 | Tripeptidyl-peptidase 1 |
| AIPGENE4319 | 21 | 9.3 | 2.2 | 3.0 | Arylsulfatase B |
| AIPGENE27525 | 40 | 43 | 1.6 | 3.0 | Betaine--homocysteine S-methyltransferase 1 |
| AIPGENE20697 | 40 | 56 | 1.0 | 2.9 | Betaine--homocysteine S-methyltransferase 1 |
| AIPGENE25614 | 10 | 11 | 2.1 | 2.6 | Lysosome membrane protein 2 |
| AIPGENE7709 | 29 | 18 | 2.8 | 2.4 | E3 ubiquitin-protein ligase MIB1 |
| AIPGENE14803 | 28 | 15 | 2.7 | 2.3 | E3 ubiquitin-protein ligase MIB1 |
| AIPGENE462 | 11 | 8.5 | 1.2 | 2.2 | Chymotrypsinogen B |
| AIPGENE28006 | 30 | 22 | 2.2 | 2.0 | Alpha-N-acetylglucosaminidase |
| AIPGENE19389 | 29 | 25 | 2.1 | 1.8 | N-acylethanolamine-hydrolyzing acid amidase |
| AIPGENE28710 | 41 | 39 | 2.4 | 1.7 | Arylsulfatase B |
| AIPGENE20962 | 110 | 89 | 1.5 | 1.5 | Lysosomal-trafficking regulator |
| AIPGENE28230 | 24 | 5.4 | 3.7 | 1.3 | Sulfotransferase 1C2A |

^a^ Ordered by SymGas/SymEpi.

^b^ Genes potentially involved in protein degradation are highlighted.

^c^ Numbers are rounded to two significant digits.

**Table S9. Module 5 genes involved in cnidocyte function**

| **Gene ID**^a,b^ | **Average expression level (TPM)**^c^ | | **Fold change**^c^ | | **Annotation** |
| --- | --- | --- | --- | --- | --- |
|  | **ApoEpi** | **SymEpi** | **ApoEpi/ApoGas** | **SymEpi/SymGas** |  |
| AIPGENE23882 | 10 | 9.0 | 5.7 | 90 | Nematocyte expressed protein 6 |
| AIPGENE21874 | 11 | 35 | 11 | 19 | Equinatoxin-2 |
| AIPGENE3089 | 15 | 39 | 3.8 | 15 | Toxin PsTX-60B |
| AIPGENE8715 | 12 | 37 | 8.0 | 15 | Toxin CfTX-2 |
| AIPGENE5924 | 63 | 320 | 2.6 | 10 | minicollagen 1 |
| AIPGENE3166 | 19 | 44 | 6.3 | 9.8 | Toxin AvTX-60A |
| AIPGENE11611 | 58 | 190 | 2.9 | 8.1 | Nematocyst outer wall antigen |
| AIPGENE4696 | 45 | 69 | 37 | 8.1 | Phospholipase A2 |
| AIPGENE13265 | 46 | 76 | 2.9 | 6.3 | Nematocyte expressed protein 6 |
| AIPGENE17670 | 9.6 | 30 | 5.8 | 4.3 | Toxin CaTX-A |
| AIPGENE3053 | 15 | 13 | 2.4 | 4.2 | Toxin AvTX-60A |
| AIPGENE17675 | 8.5 | 47 | 2.2 | 3.7 | Toxin CaTX-A |
| AIPGENE19223 | 6.3 | 6.8 | 2.8 | 3.4 | Toxin CaTX-A |

^a^ Ordered by SymEpi/SymGas.

^b^ Four previously well characterized nematocyte markers are highlighted; two of them are shown in Fig. 1G.

^c^ Numbers are rounded to two significant digits.

**Table S10. Module 5 genes involved in responses to mechanical stimuli**

| **Gene ID**^a,b^ | **Average expression level (TPM)**^c^ | | **Fold change**^c^ | | **Annotation** |
| --- | --- | --- | --- | --- | --- |
|  | **ApoEpi** | **SymEpi** | **ApoEpi/ApoGas** | **SymEpi/SymGas** |  |
| AIPGENE6717 | 20 | 55 | 2.8 | 15 | Amyloid beta A4 |
| AIPGENE26233 | 54 | 67 | 3.2 | 7.5 | Tetraspan membrane protein of hair cell stereocilia |
| AIPGENE7279 | 130 | 190 | 6.9 | 7.2 | Nephrocystin-3 |
| AIPGENE727 | 21 | 13 | 3.6 | 6.9 | Acid-sensing ion channel 3 |
| AIPGENE13869 | 48 | 300 | 3.0 | 6.2 | YadA domain protein |
| AIPGENE16071 | 15 | 14 | 5.2 | 5.2 | Acid-sensing ion channel 3 |
| AIPGENE7259 | 58 | 58 | 5.1 | 4.8 | Nephrocystin-3 |
| AIPGENE7280 | 31 | 43 | 3.5 | 4.7 | Nephrocystin-3 |
| AIPGENE7264 | 150 | 120 | 2.5 | 4.4 | Nephrocystin-3 |
| AIPGENE10398 | 44 | 40 | 4.7 | 4.1 | Nephrocystin-3 |
| AIPGENE18522 | 12 | 11 | 3.3 | 3.7 | Acid-sensing ion channel 3 |
| AIPGENE7455 | 7.1 | 6.5 | 3.2 | 3.6 | Acid-sensing ion channel 3 |
| AIPGENE565 | 44 | 12 | 6.6 | 3.2 | Transient receptor potential cation channel subfamily A member 1 TRPA1 |
| AIPGENE3114 | 20 | 28 | 5.5 | 3.2 | Nephrocystin-3 |
| AIPGENE3223 | 78 | 75 | 8.1 | 3.0 | Nephrocystin-3 |
| AIPGENE13728 | 29 | 26 | 5.8 | 2.7 | Transient receptor potential cation channel subfamily A member 1 TRPA1 |
| AIPGENE4254 | 43 | 54 | 2.5 | 2.3 | Nephrocystin-3 |
| AIPGENE1377 | 34 | 32 | 1.2 | 2.0 | Acid-sensing ion channel 3 |
| AIPGENE15296 | 20 | 18 | 2.5 | 2.0 | Nephrocystin-3 |
| AIPGENE20487 | 350 | 270 | 1.9 | 1.3 | Nephrocystin-3 |
| AIPGENE20323 | 79 | 59 | 2.0 | 1.2 | Nephrocystin-3 |

^a^ Ordered by SymEpi/SymGas.

^b^ Genes potentially required for normal ciliary development and function are highlighted.

^c^ Numbers are rounded to two significant digits.

**Table S11. Potential cnidocyte marker genes in Cluster 11**

| **Gene ID**^a,b^ | **Fold change**^c^ | **Adjusted *p* value** | **Annotation** |
| --- | --- | --- | --- |
| AIPGENE5924 | 37 | 5E-25 | Minicollagen 1 |
| AIPGENE23847 | 20 | 3E-18 | Minicollagen 2 |
| AIPGENE25615 | 14 | 7E-40 | Venom prothrombin activator omicarin-C |
| AIPGENE11170 | 7.4 | 3E-12 | Latrophilin-3 |
| AIPGENE11611 | 6.1 | 9E-33 | Nematocyst outer wall antigen |
| AIPGENE29070 | 5.5 | 1E-05 | Matrilin-2 |
| AIPGENE25580 | 5.0 | 9E-88 | Lactadherin OS=Rattus norvegicus |
| AIPGENE2293 | 3.7 | 9E-88 | D-galactoside-specific lectin |
| AIPGENE23596 | 2.0 | 6E-49 | L-rhamnose-binding lectin CSL1 |

^a^ Ordered by Fold change.

^b^ Genes shown in Fig. 1G are highlighted.

^c^ Fold-changes calculated by comparing the cells in Cluster 11 with all the other cells.

**Table S12. Potential gastrodermal-cell marker genes in Cluster 2, 7, and/or 10**

| **Cluster** | **Gene ID**^a,b^ | **Fold change**^c^ | **Adjusted *p* value** | **Annotation** |
| --- | --- | --- | --- | --- |
| 2 | AIPGENE21625 | 2.5 | 1E-18 | Titin |
| 2 | AIPGENE3988 | 2.0 | 4E-05 | Tropomyosin |
| 7 | AIPGENE6165 | 8.2 | 5E-22 | Integrin alpha-8 |
| 7 | AIPGENE23790 | 6.0 | 3E-60 | Integrin beta-6 |
| 7 | AIPGENE27486 | 5.5 | 4E-21 | Tropomyosin alpha-4 |
| 7 | AIPGENE23262 | 4.5 | 6E-56 | Fibroblast growth factor receptor 1 |
| 7 | AIPGENE5861 | 4.5 | 2E-18 | Integrin beta-2 |
| 7 | AIPGENE22443 | 3.7 | 1E-28 | Soma ferritin |
| 7 | AIPGENE21255 | 3.7 | 2E-12 | Integrin alpha-4 |
| 7 | AIPGENE16019 | 3.0 | 4E-22 | Fibroblast growth factor receptor 3 |
| 7 | AIPGENE8440 | 2.7 | 1E-11 | Integrin alpha-6 |
| 7 | AIPGENE15963 | 2.5 | 1E-11 | Laminin beta-1 |
| 7 | AIPGENE17012 | 2.2 | 3E-12 | Laminin alpha-4 |
| 7 | AIPGENE6003 | 2.0 | 4E-07 | Cathepsin L |
| 7 | AIPGENE26156 | 2.0 | 5E-17 | Cathepsin L |
| 7 | AIPGENE2368 | 2.0 | 5E-06 | Laminin gamma-1 |
| 10 | AIPGENE3975 | 5.5 | 1E-25 | Fibroblast growth factor receptor |
| 10 | AIPGENE22204 | 3.7 | 3E-23 | Cathepsin B |
| 10 | AIPGENE23262 | 2.5 | 7E-11 | Fibroblast growth factor receptor 1 |
| 10 | AIPGENE26157 | 2.2 | 7E-13 | Cathepsin L |

^a^ Ordered by Fold change within each cluster.

^b^ Potential marker genes were selected according to cell markers identified in other cnidarian species.

^c^ Fold-changes calculated by comparing the cells in Cluster 2, 7, or 10 with all the other cells.

**Table S13. Cluster 10 marker genes associated with lysosomal organization and function**

| **Gene ID**^a^ | **Fold change**^b^ | **Adjusted *p* value** | **Annotation** |
| --- | --- | --- | --- |
| AIPGENE22204 | 3.7 | 3E-23 | Cathepsin B |
| AIPGENE21470 | 3.3 | 3E-16 | Proactivator polypeptide |
| AIPGENE5531 | 3.3 | 3E-21 | Niemann-Pick C1 NPC1 |
| AIPGENE721 | 3.0 | 6E-11 | Sialate O-acetylesterase |
| AIPGENE19216 | 2.5 | 2E-23 | Acid ceramidase |
| AIPGENE1519 | 2.5 | 5E-19 | Tyrosine-protein kinase SRC42A |
| AIPGENE9971 | 2.5 | 2E-19 | Lysosomal aspartic protease |
| AIPGENE26157 | 2.2 | 7E-13 | Cathepsin L |
| AIPGENE22539 | 2.2 | 1E-05 | Epididymal secretory protein NPC2 |
| AIPGENE3899 | 2.0 | 2E-11 | CD63 antigen |
| AIPGENE756 | 2.0 | 4E-30 | Putative phospholipase B-like 2 |
| AIPGENE5532 | 2.0 | 3E-07 | Niemann-Pick C1 NPC1 |
| AIPGENE18432 | 2.0 | 4E-21 | Low-density lipoprotein receptor-related protein 6 |

^a^ Ordered by Fold change.

^b^ Fold-changes calculated by comparing the cells in Cluster 10 with all the other cells.

**Table S14. Cluster 10 marker genes associated with glucose transport and its positive regulation**

| **Gene ID**^a,b^ | **Fold change**^c^ | **Adjusted *p* value** | **Annotation** |
| --- | --- | --- | --- |
| AIPGENE12082 | 1.8 | 4E-24 | Solute carrier family 2, facilitated glucose transporter member 1 |
| AIPGENE10547 | 1.5 | 2E-25 | C2 domain-containing protein 5 |
| AIPGENE10700 | 1.3 | 4E-04 | Bifunctional protein NCOAT |

^a^ Ordered by Fold change.

^b^ Cyan highlight, the gene encoding glucose transporter AipGLUT1α, as investigated further in this study.

^c^ Fold-changes calculated by comparing the cells in Cluster 10 with all the other cells.

**Table S15. Cluster 10 marker genes associated with cholesterol transport and homeostasis**

| **Gene ID**^a^ | **Fold change**^b^ | **Adjusted *p* value** | **Annotation** |
| --- | --- | --- | --- |
| AIPGENE5531 | 3.3 | 3E-21 | Niemann-Pick C1 NPC1 |
| AIPGENE22539 | 2.2 | 9E-06 | Epididymal secretory protein NPC2 |
| AIPGENE5532 | 2.0 | 3E-07 | Niemann-Pick C1 NPC1 |
| AIPGENE18432 | 2.0 | 4E-21 | Low-density lipoprotein receptor-related protein 6 |

^a^ Ordered by Fold change.

^b^ Fold-changes calculated by comparing the cells in Cluster 10 with all the other cells.

**Table S16. Cluster 7 marker genes associated with extracellular matrix, cell adhesion, and cell-cell signaling**

| **Gene ID^a^** | **Fold change^b^** | **Adjusted *p* value** | **Annotation** |
| --- | --- | --- | --- |
| AIPGENE10422 | 18 | 1E-85 | Collagen alpha-1(IV) chain |
| AIPGENE10546 | 11 | 4E-65 | Collagen alpha-2(IV) chain |
| AIPGENE8588 | 9.0 | 2E-60 | Testican-2 |
| AIPGENE1266 | 8.2 | 6E-28 | Somatomedin-B and thrombospondin type-1 domain-containing protein |
| AIPGENE6165 | 8.2 | 5E-22 | Integrin alpha-8 |
| AIPGENE27343 | 7.4 | 2E-46 | Collagen alpha-4(VI) chain |
| AIPGENE23790 | 6.0 | 3E-60 | Integrin beta-6 |
| AIPGENE2489 | 6.0 | 6E-34 | Matrilin-2 |
| AIPGENE3595 | 5.5 | 1E-32 | Serotransferrin |
| AIPGENE9466 | 5.5 | 8E-16 | Fibroblast growth factor 18 |
| AIPGENE7597 | 5.0 | 7E-46 | Zonadhesin |
| AIPGENE1737 | 5.0 | 6E-44 | Protein Wnt |
| AIPGENE23312 | 5.0 | 1E-39 | Hemicentin-1 |
| AIPGENE12342 | 5.0 | 4E-28 | Collagen alpha-3(VI) chain |
| AIPGENE22492 | 5.0 | 7E-37 | Glypican-6 |
| AIPGENE7897 | 5.0 | 2E-15 | Hemicentin-1 |
| AIPGENE23262 | 4.5 | 6E-56 | Fibroblast growth factor receptor 1 |
| AIPGENE17918 | 4.5 | 2E-16 | Hemicentin-2 |
| AIPGENE10250 | 4.5 | 6E-21 | Collagen alpha-6(VI) chain |
| AIPGENE20628 | 4.5 | 6E-47 | Collagen alpha-4(VI) chain |
| AIPGENE23383 | 4.1 | 5E-38 | Hemicentin-2 |
| AIPGENE29129 | 4.1 | 6E-40 | Periostin |
| AIPGENE6975 | 4.1 | 8E-43 | Neurofascin |
| AIPGENE16436 | 4.1 | 5E-53 | Protocadherin Fat 1 |
| AIPGENE27342 | 4.1 | 2E-07 | Agrin |
| AIPGENE11184 | 4.1 | 3E-16 | Collagen alpha-1(VII) chain |
| AIPGENE17222 | 3.7 | 3E-29 | Agrin |
| AIPGENE25153 | 3.7 | 6E-24 | Collagen alpha-5(VI) chain |
| AIPGENE10561 | 3.7 | 4E-17 | Collagen alpha-1(XXVI) chain |
| AIPGENE21255 | 3.7 | 2E-12 | Integrin alpha-4 |
| AIPGENE12362 | 3.3 | 8E-08 | Collagen alpha-6(VI) chain |
| AIPGENE20627 | 3.3 | 3E-09 | Collagen alpha-3(VI) chain |
| AIPGENE7612 | 3.3 | 2E-26 | Reticulocyte-binding protein 2 homolog a |
| AIPGENE8011 | 3.0 | 5E-15 | SCO-spondin |
| AIPGENE16749 | 3.0 | 3E-11 | Talin-2 |
| AIPGENE18950 | 3.0 | 1E-06 | Metastasis suppressor protein 1 |
| AIPGENE3608 | 3.0 | 3E-07 | Receptor-type tyrosine-protein phosphatase delta |
| AIPGENE12347 | 3.0 | 7E-15 | Collagen alpha-6(VI) chain |
| AIPGENE266 | 3.0 | 1E-10 | Dystroglycan |
| AIPGENE23512 | 3.0 | 1E-06 | Protein Wnt-4 |
| AIPGENE3034 | 2.7 | 2E-24 | Mammalian ependymin-related protein 1 |
| AIPGENE8440 | 2.7 | 1E-11 | Integrin alpha-6 |
| AIPGENE29194 | 2.7 | 3E-14 | Catenin delta-2 |
| AIPGENE4075 | 2.5 | 4E-14 | Proprotein convertase subtilisin/kexin type 5 |
| AIPGENE15963 | 2.5 | 1E-11 | Laminin subunit beta-1 |
| AIPGENE14892 | 2.5 | 9E-16 | Calsyntenin-2 |
| AIPGENE26184 | 2.5 | 7E-11 | Fibrillin-1 |
| AIPGENE9673 | 2.5 | 6E-10 | Fibulin-1 |
| AIPGENE22521 | 2.2 | 9E-10 | Glypican-1 |
| AIPGENE25192 | 2.2 | 8E-22 | Collagen alpha-1(XXIV) chain |
| AIPGENE15522 | 2.2 | 2E-06 | Protocadherin Fat 1 |
| AIPGENE17012 | 2.2 | 3E-12 | Laminin subunit alpha-4 |
| AIPGENE19606 | 2.2 | 2E-05 | Thrombospondin-1 |
| AIPGENE18534 | 2.2 | 1E-20 | Collagen alpha-2(I) chain |
| AIPGENE5847 | 2.2 | 8E-07 | Semaphorin-5A |
| AIPGENE25893 | 2.2 | 3E-11 | Collagen alpha-1(XXVII) chain B |
| AIPGENE8276 | 2.0 | 2E-08 | Myosin-10 |
| AIPGENE18348 | 2.0 | 4E-05 | Fibrillin-1 |
| AIPGENE8376 | 2.0 | 6E-12 | Collagen alpha-2(I) chain |
| AIPGENE2639 | 2.0 | 5E-06 | Afadin |
| AIPGENE6961 | 2.0 | 2E-10 | Neurofascin |
| AIPGENE2368 | 2.0 | 5E-06 | Laminin subunit gamma-1 |

^a^ Ordered by Fold change.

^b^ Fold-changes calculated by comparing the cells in Cluster 7 with all the other cells.

**Table S17. Expression patterns of the five major glucose and ammonium transporters**

| **Gene**^a^ | **Average expression level (TPM)**^b^ | | **Fold change**^b^ | |
| --- | --- | --- | --- | --- |
|  | **SymEpi** | **SymGas** | **SymEpi/ApoEpi** | **SymGas/ApoGas** |
| AipSLC2A8α (AIPGENE2706) | 2.2 | 21 | 0.84 | 4.7 |
| AipGLUT1α (AIPGENE12082) | 33 | 174 | 0.67 | 1.0 |
| AipGLUT8α (AIPGENE18406) | 8.9 | 70 | 2.1 | 6.8 |
| AipRhBG1(AIPGENE18105) | 20 | 410 | 6.8 | 170 |
| AipAMT1 (AIPGENE17420) | 98 | 25 | 1.1 | 1.4 |

^a^ Genes are listed in the order in which they are shown in Fig. 2 A, B, and E-S.

^b^ Numbers are rounded to two significant digits.

**Table S18. Peptides used as antigens to generate custom antibodies**

| **Target ID** | **Peptide sequence**^a,b^ |
| --- | --- |
| AipAMT1 (AIPGENE17420) | NGGSQGSIASKGDA |
| AipRhBG1 (AIPGENE18105) | IDQESGHSINSIKS |
| AipSLC2A8α (AIPGENE2706) | ALLGGPLGGWLIEAFGRKGG |
| AipGLUT1α (AIPGENE12082) | NSPEKVIKDYYKKYGHEFTD |
| AipGLUT8α (AIPGENE18406) | PDMDIDFEIA |

^a^ Peptides were selected from non-transmembrane regions with high predicted antigenicity and hydrophilicity.

^b^ An additional cysteine was added to either the N- or C-terminus of each peptide to facilitate its conjugation to keyhole-limpet hemocyanin to create the antigens used for immunization.

**Table S19. Primers used to amplify the transporter genes**

| **Target ID** | **Forward primer** | **Reverse primer** |
| --- | --- | --- |
| AipAMT1  (AIPGENE17420) | atgatgaacgtaactacagaacagac | ttacaatttttcatcaagagctccgtg |
| AipRhBG1  (AIPGENE18105) | atgtgctcggcaatcataacaacacgc | ttaatcactcttgatggaattgatggaatgtcc |
| AipSLC2A8α  (AIPGENE2706) | GCTAGCatggccaatcaaaatattcaaagtagc | GGATCCttatatgcgttcatattcag |
| AipGLUT1α  (AIPGENE12082) | GCTAGCatggaggaggaaaagggggagac | GGATCCttaatcatcgtccccaacagaac |
| AipGLUT8α  (AIPGENE18406) | GCTAGCatgaatcgttcattactcaagcg | GGATCCttaaagtctgctttctacctctgc |

Sequences in color indicate the enzyme-cutting sites for *Nhe*I and *Bam*HI, used in cloning the amplified products into plasmid pSH100 (see text). Gene-specific-primer sequences are in lower case. No enzyme-cutting sites were added for the ammonium-transporter genes, as they were cloned using a TA-cloning protocol.

**Data S1.**

Tissue-specific transcriptomic profiles of Aiptasia in different symbiotic states

**Data S2.**

Differentially expressed genes identified from multi-factor analysis.

**Data S3.**

Potential cell-type markers identified in single-cell RNA-seq.

**Data S4.**

Gene-set-enrichment analysis of identified Cluster 10 marker genes.

**Data S5.**

Gene-set-enrichment analysis of identified Cluster 7 marker genes.
